# Phylogenomics of the leaf-footed bug subfamily Coreinae (Hemiptera: Coreidae): applicability of ultraconserved elements at shallower depths

**DOI:** 10.1101/2020.03.18.997569

**Authors:** Michael Forthman, Christine W. Miller, Rebecca T. Kimball

## Abstract

Baits targeting invertebrate ultraconserved elements (UCEs) are becoming more common for phylogenetic studies. Recent studies have shown that invertebrate UCEs typically encode proteins — and thus, are functionally different from more conserved vertebrate UCEs —can resolve deep divergences (e.g., superorder to family ranks). However, the ability of the invertebrate UCE baits to robustly resolve relationships at shallower phylogenetic scales (i.e., tribes and congeners) has been generally limited to Coleoptera and Hymenoptera. Here, we assessed the ability of a recently designed Hemiptera UCE bait set to reconstruct more recent phylogenetic relationships in the largest leaf-footed bug subfamily, the Coreinae (Hemiptera: Coreidae), using a taxon-rich sample representing 21 of the 32 coreine tribes. Many well-supported, novel relationships were congruent in maximum likelihood and summary coalescent analyses. We also found evidence for the para- and polyphyly of several tribes and genera of Coreinae, as well as the subfamilies Coreinae and Meropachyinae. Our study, along with other recent UCE studies, provides evidence that UCEs can produce robust and novel phylogenetic hypotheses at various scales in invertebrates. Additionally, we used different DNA extraction and target enrichment protocols and recovered more UCE data using a touch-down hybridization approach.

## Introduction

Next-generation sequencing (NGS) technology has made the generation of thousands of orthologous loci throughout the genome achievable for many non-model organisms. For molecular phylogenetics, one of the advantages of such data is the potential to resolve challenging nodes in the Tree of Life across various temporal scales (e.g., Faircloth et al. 2012, Lemmon et al. 2012, Li et al. 2013). Ultraconserved elements (UCEs) are one such class of loci that can be obtained by using target capture approaches and NGS (see Faircloth et al. 2012). These loci have been widely used in phylogenetic estimation since Faircloth et al. (2012) introduced their utility in anchoring loci for phylogenomic analysis in vertebrates. In vertebrates, UCEs are highly conserved regions of the genome that are believed to be primarily non-coding regulators of gene expression (Bejerano et al. 2004, Sandelin et al. 2004, Woolfe et al. 2004, Pennacchio et al. 2006). The high conservation of UCEs across divergent taxa (e.g., >90% conserved across amniotes) (Bejerano et al. 2004, Faircloth et al. 2012) has allowed capture of sequences that can resolve deep divergences using a single set of baits (e.g., Crawford et al. 2012, McCormack et al. 2012, Faircloth et al. 2013, Gilbert et al. 2015). Additionally, as sequence variability increases away from the conserved core region (i.e., flanking regions), more recent divergences between species and populations can also be achieved (e.g., Smith et al. 2014, Manthey et al. 2016).

Although the use of UCEs in invertebrates is conceptually similar as it is applied in vertebrates, baits have been designed to target genomic regions with more liberal thresholds of conservation across taxa (Faircloth 2017), often requiring multiple baits for the same region to maximize capture of divergent taxa. Furthermore, recent empirical tests of UCE baits have shown that these loci are primarily protein-coding in invertebrates (Bossert and Danforth 2018, Kieran et al. 2019) making invertebrate UCEs fundamentally different from those found in vertebrates. Regardless, invertebrate UCE bait sets have be shown to resolve deep divergences (e.g., superorder to family ranks) in several taxonomic groups (e.g., Baca et al. 2017, Starrett et al. 2017, Van Dam et al. 2017, Kieran et al. 2019). However, there have been few studies demonstrating the utility of these primarily protein-coding UCEs at more shallow evolutionary timescales (i.e., subfamily to congeners) for taxa in which bait sets are available (e.g., Van Dam et al. 2017, Hedin et al. 2018, Bossert et al. 2019). Recently, Kieran et al. (2019) empirically tested a UCE bait set for the insect order Hemiptera (Faircloth 2017), showing its utility in resolving phylogenetic relationships among the suborders down to intrafamilial relationships in Reduviidae and Coreidae with a small sample of taxa. The utility of UCEs in hemipteran phylogenetics has also been shown at a relatively shallower scale as seen in Forthman et al. (2019), who focused on interfamilial and intersubfamilial relationships in the superfamily Coreoidea. However, the ability of the invertebrate UCE bait sets to robustly resolve phylogenetic relationships among tribes and congeners has not been widely demonstrated beyond a few studies primarily focused on Coleoptera and Hymenoptera (e.g., Van Dam et al. 2017, Ješovnik et al. 2017, Bossert et al. 2019, Branstetter and Longino 2019).

Leaf-footed bugs, or Coreidae (Hemiptera: Heteroptera), are a charismatic group of phytophagous insects (Fig. 1) that also includes several pests of agricultural systems (see Gentry 1965, Nonveiller 1984, Mitchell 2000). With 2,571 extant species described in four subfamilies and 37 tribes, this is the largest family of the Coreoidea (CoreoideaSF Team 2019). The worldwide Coreinae is by far the largest coreid subfamily with 2,320 (90%) species (372 genera, 32 tribes). Some of the largest, stoutest terrestrial heteropterans are members of this subfamily (e.g., species of *Pachylis*, *Thasus*, and *Petascelis*; Schuh and Slater 1995, Fernandes et al. 2015), but body forms also vary from sticklike (e.g., *Tylocryptus*, *Prionotylus*) to extravagant foliaceous or spined expansions (e.g., species of the tribe Phyllomorphini). While many species are dull in appearance, some are brightly colored and iridescent (e.g., *Petalops*, *Diactor*, *Phthiadema*). The hind legs of males in many species are known to be sexually selected weapons that exhibit variation in size, shape, and armature (Eberhard 1998, Emlen 2008, Okada et al. 2011, Procter et al. 2012). Fighting behaviors are also variable in species that exhibit male-male competition; e.g., some species grapple end-to-end (e.g., *Narnia femorata*; Nolen et al. 2017), while others kick, flip, and squeeze one another face-to-face (e.g., *Mictis profana*; (Tatarnic and Spence 2013). Aside from their diverse morphology, Coreinae are well known for their odious alarm pheromones (Aldrich and Blum 1978, Leal and Kadosawa 1992), paternal care in *Phyllomorpha* (e.g., García-González et al. 2003), and gregariousness in nymphs (e.g., Aldrich and Blum 1978, Flanagan 1994, Miyatake 1995). Furthermore, species of the genus *Holhymenia* superficially appear to be wasp mimics (Pereira et al. 2013). Given these captivating morphological and behavioral features, the subfamily Coreinae offers an excellent opportunity to investigate the evolution of various traits and their possible correlates. However, a well-resolved, taxon-rich phylogeny of Coreinae is lacking. Thus, a comprehensive, robust phylogeny of the group is first needed before evolutionary questions can be investigated.

**Figure 1.**
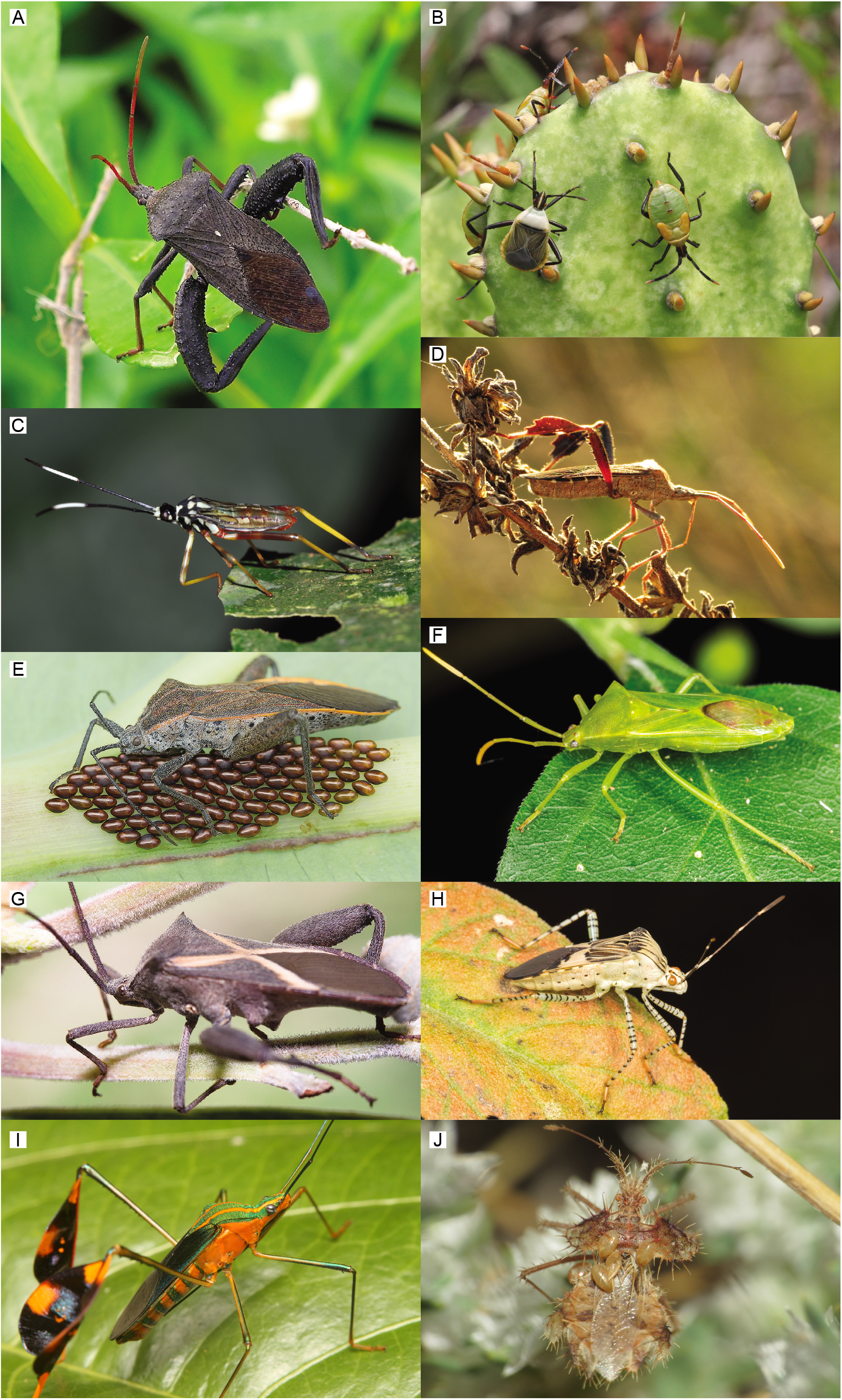
Images of representative Coreinae. (a) *Acanthocephala femorata* (Acanthocephalini) (© 2015 Kala King). (b) *Chelinidea vittiger* (Chelinideini) (© 2017 Mary Keim). (c) *Holhymenia* sp. nymph (Anisoscelini) (© 2011 Arthur Anker). (d) *Leptoglossus* sp. (Anisoscelini) (© 2006 Sean McCann). (e) *Physomerus grossipes* (Acanthocorini) (© 2014 Anthony Kei C Wong). (f) *Savius jurgiosus* (Discogastrini) (© 2016 Jeff Gruber). (g) *Mictis profana* (Mictini) (© 2011 Jon Clark). (h) *Hypselonotus* sp. (Hypselonotini) (© 2015 Jeff Gruber). (i) *Diactor* sp. (Anisoscelini) (© 2015 Jorge Restrepo). (j) *Phyllomorpha laciniata* (Phyllomorphini) (© 2015 Simon Oliver).

Although a few molecular phylogenetic analyses of the Coreidae have been performed (Fang and Nie 2007, Pan et al. 2007, Pan et al. 2008), the most comprehensive (with respect to taxon sampling) investigation of coreine phylogeny comes from Li’s (1997) morphological phylogenetic analysis of the family Coreidae. Li (1997) recovered a paraphyletic Coreinae with respect to Meropachyinae, which has also been supported by Li (1996), Kieran et al. (2019), and Forthman et al. (2019). Li’s (1997) analysis further suggested the tribe Colpurini to be the earliest diverging lineage within the Coreinae, which was also supported by Li (1996). Although Li (1997) found evidence that at some coreine tribes are not monophyletic, many of the 21 sampled coreine tribes were represented by a single species. Furthermore, many morphological traits analyzed by Li (1997) were recently shown to exhibit homoplasy (Forthman et al. 2019). Since Li’s (1997) analysis, hypotheses about inter-and intratribal relationships within the Coreinae remain to be investigated more comprehensively.

Here, we use UCEs to develop a more comprehensive understanding of the phylogeny of Coreinae and assess its utility in reconstructing relationships at shallow evolutionary timescales in this subfamily. Our taxon sampling includes representatives of 21 morphologically diverse tribes of Coreinae (out of 32), which is the most extensive sampling of tribes since Li’s (1997) analysis. This included several of the most and least speciose tribes (e.g., Hypselonotini and Phyllomorphini, respectively) from all major biogeographic regions. With our taxon sampling, we first examine the circumscription of currently recognized tribes, such as the Anisoscelini and Coreini whose circumscription has varied among studies. Secondly, we test previous hypotheses regarding phylogenetic relationships within Coreinae, such as the early divergence of Colpurini from other tribes (Schaefer 1965, Ahmad 1970), the paraphyly of Nematopodini (Kieran et al. 2019, Forthman et al. 2019), and the non-monophyly of the subfamily with respect to the Meropachyinae (Li 1996, 1997, Kieran et al. 2019, Forthman et al. 2019). Third, we explore the suitability of UCEs at shallower scales by including multiple species within several genera whose limits have been generally uncontroversial. Lastly, previous studies have reported improvements in locus recovery by implementing alternative approaches to molecular protocols, such as target enrichment (e.g., Li et al. 2013, Paijmans et al. 2016) or DNA extraction protocols (e.g., Chen et al. 2010) that can impact that quality and quantity of samples — including dried museum samples — for downstream processing. Thus, we examine the use of different DNA extraction and target enrichment approaches and compare overall locus recovery among them.

## Material and methods

### Taxon sampling

A total of 124 taxa were sampled for this study, including 104 species of Coreinae from 21 tribes. For 25 of our taxa, we obtained contigs from (Kieran et al. 2019) (NCBI Sequence Read Archive SRP161492) and (Forthman et al. 2019) (NCBI BioProject PRJNA531965). We generated new UCE data for the remaining taxa (Table S1) following the protocols discussed below.

### DNA extraction

For all new data, genomic DNA was extracted using a 1) Gentra Puregene Tissue, 2) Qiagen DNeasy Blood and Tissue kit (hereafter DNeasy), or 3) Qiagen DNeasy Blood and Tissue kit coupled with Qiagen QIAquick PCR purification kit (hereafter DNQIA; see Knyshov et al. 2019). Depending on the size of specimens, we used any part of the body (legs, abdomen, thorax, head) or the entire body from ethanol-preserved, silica-bead preserved, frozen, or dried specimens. For the Puregene Tissue Kit, we followed the manufacturer’s protocol for 5–10 mg tissue with several modifications: 10 μL of proteinase K was added to samples and incubated for 24–48 hours; samples were incubated at 37°C for 60 mins after adding 1.5 μL of RNase A solution, after which samples were placed on ice for 3 mins; two centrifugations, with ice incubation for 5 mins in between, were performed to ensure precipitated proteins formed a tight pellet; 600 μL of 100% ethanol was used for the first wash and centrifuged for 10 mins; and 50– 100 μL of molecular grade water or Puregene DNA Hydration Solution was used to resuspend isolated DNA. We also followed the manufacturer’s protocol for the DNeasy kit, but with fewer modifications: tissue was incubated in 180–190 μL Buffer ATL and 10–20 μL proteinase K for 24–48 hours (200 μL total solution volume), and DNA eluted once or twice with 50 μL Buffer AE depending on the size of the specimens we extracted from.

To assess which extraction protocol may perform better with dried museum material that is likely dominated by degraded DNA, we either used the DNeasy protocol described above or a modified version of it that follows Knyshov et al. (2019) (i.e., DNQIA). The latter protocol is specifically designed to extract degraded DNA >100 bp in length. Briefly, the DNQIA protocol follows the DNeasy kit up to the first centrifugation, using a QIAquick spin column. The samples are then subjected to the Qiagen QIAquick PCR purification kit, replacing AW1 and AW2 washes with PE buffer. Under the DNQIA protocol, samples are eluted in 30 μL EB buffer. We initially attempted to extract genomic DNA from six specimens using the DNeasy protocol (collected 1946–2016) and ten others using the DNQIA protocol (collected 1935–2015). Two museum samples (collected 1980 and 1987) initially subjected to the DNeasy protocol did not yield extracts of sufficient concentration nor produce visual bands after gel electrophoresis (see below for details on methods); these samples were re-extracted using the DNQIA protocol.

DNA quality was assessed with 1% agarose gel electrophoresis and quantified with a Qubit 2.0 fluorometer. Samples were then normalized to 10–20 ng/μL, and high molecular weight samples were fragmented into 200–1000 bp using a Biorupter UCD-300 sonication device (4–10 cycles of 30 s on/30 s) or a Covaris M220 Focused-ultrasonicator (20–60 s).

### Library construction, target enrichment, and sequencing

We constructed libraries using the modified KAPA Hyper Prep Kit protocol of Forthman et al. (2019). Briefly, half volume reactions were used for all steps, as well as iTru universal adapter stubs and 8 bp dual-indexes (Glenn et al. 2016). Library amplification involved initial denaturation at 98°C for 3 min, followed by 14 cycles of 98°C for 30 s, 60°C for 30 s, and 72°C for 30 s, with a final extension at 72°C for 5 min. Amplified libraries were inspected by gel electrophoresis, quantified with Qubit, combined into 1000 ng pools using equimolar amounts, dried at 60°C, and resuspended in 14 μL IDTE.

For target enrichment, we used a custom myBaits kit, based on Faircloth’s (2017) probe set that was subsampled by Forthman et al. (2019). For some samples, we followed Forthman et al.’s (2019) enrichment protocol while others were subjected to a modified version of the protocol. We refer to the protocols as TE and TE-touchdown, respectively. For our TE protocol, a hybridization mixture with half volume of baits (2.75 μL) and 2.75 μL molecular-grade water was hybridized with each library pool at 65°C for 16–24 hours. In the TE-touchdown protocol, baits were to hybridize with library pools at 65°C for 18 hours followed by 18 hours at 62°C. However, samples were prematurely removed during the TE-touchdown protocol before hybridization was complete. At the recommendation of Arbor Biosciences, we added an additional 2.75 μL baits to these samples and re-ran the hybridization protocol to completion; additional baits were added given that the initial beads were subjected to the 95°C denaturation step, which may limit their effectiveness during hybridization per Arbor Biosciences.

Dynabeads M-280 Streptavidin beads were then bound to bait-target hybrids, washed four times at 65°C (TE) or 62°C (TE-touchdown) and resuspended in 30 μL IDTE. For the post-capture PCR amplification mix, 2.5 μL each of 5 μM iTru P5/P7 primers (Glenn et al. 2016) were added. We performed 14–17 cycles of post-capture amplification following manufacturer’s protocol, except we used an annealing temperature of 65°C (TE) or 62°C (TE-touchdown) and an extension period of 45 s. Hydrophobic Sera-Mag SpeedBeads Carboxyl Magnetic Beads were used for post-amplification cleanup, followed by two washes in freshly prepared 70% ethanol and resuspension in 22 μL IDTE. We quantified enriched library pools with Qubit, pooled all library pools in equimolar amounts, and sequenced on a single Illumina HiSeq3000 lane (2×100) at the University of Florida’s Interdisciplinary Center for Biotechnology Research (ICBR).

### Sequence data processing and alignment

Sequence data were processed following Forthman et al. (2019). Briefly, sequence reads were demultiplexed at the sequencing facility, and adapters were trimmed with illumiprocessor (Faircloth 2013, Bolger et al. 2014). Duplicate reads were filtered using PRINSEQ-lite v0.20.4 (Schmieder and Edwards 2011), and the remaining reads error-corrected with QuorUM v1.1.0 (Marçais et al. 2015) and *de novo* assembled in Trinity (Grabherr et al. 2011). We identified and aligned UCE loci from our assembled contigs using PHYLUCE v1.5.0 (Faircloth 2016). Internal trimming of alignments was done with trimAl (Capella-Gutiérrez et al. 2009). Locus alignments with at least 50% and 70% of taxa (hereafter 50p and 70p, respectively) were retained for phylogenetic inferences. We also subsampled each dataset by just including the 25% most parsimony informative loci — based on raw counts — to explore the effects of this filtering strategy on phylogenetic inferences; the use of the 25% most parsimony informative loci have been shown to improve or recover similar topological support compared to estimates based on more uninformative or informative gene trees, respectively (Hosner et al. 2016, Meiklejohn et al. 2016).

### Phylogenetic estimation

For each of the four datasets, we concatenated single locus alignments in PHYLUCE and then selected the best-fit partition scheme and models of sequence evolution using PartitionFinder v2.1.1 (Lanfear et al. 2017). We used the rcluster algorithm with unlinked branch lengths and treated individual loci as separate data blocks. All models under the “raxml” option were examined (Stamatakis 2006), and the best-fit models were selected using the corrected Akaike Information Criterion (AICc) (Hurvich and Tsai 1989). Twenty partitioned maximum likelihood (ML) optimal searches were conducted in RAxML v8.2.10 (Stamatakis 2014) using random starting trees. Bootstrap (BS) support from 500 iterations were summarized on the best ML tree with SumTrees v4.0.0 (Sukumaran and Holder 2010).

We also estimated species trees from individual gene trees using an approach statistically consistent with the multispecies coalescent model (Degnan and Rosenberg 2006, 2009). We used MrAIC v1.4.6 (Nylander 2004) to select one out of the 56 models of sequence evolution for each locus alignment based on the AICc score in PhyML v3.1 (Guindon et al. 2010). We generated optimal gene trees by performing 20 ML searches in GARLI using results from MrAIC. The use of polytomous gene trees has been shown to improve species tree estimation (Zhang et al. 2017); as such, we allowed our input gene trees to have polytomies (collapsebranches = 1). One hundred bootstrap replicates were also generated by reducing the termination condition parameter by half the default value (i.e., genthreshfortopoterm = 10000) (see Zwickl 2008). Species trees were inferred from optimal gene trees in ASTRAL-III v5.6.1 (Mirarab et al. 2014, Sayyari and Mirarab 2016, Zhang et al. 2018), with clade support measured using 100 multilocus bootstrap replicates (Seo 2008).

Majority-rule consensus trees were generated in PAUP* v4.0a.16 (Swofford 2003) for the following: 1) all resolved optimal trees estimated from every analysis and 2) all optimal trees with branches having <50% bootstrap support collapsed. We then computed symmetric differences (2x Robinson-Foulds [RF]) between optimal trees (excluding outgroups) within each of these two groups to assess topological variation across our analyses and to identify if conflicting nodes still existed after poorly supported branches were collapsed.

Because our results produced a polytomy among relatively deeper branches in our majority rule consensus trees, we evaluated if the incongruence among analyses could be due to incomplete lineage sorting. Under the multispecies coalescent model, rooted three-taxon gene trees will yield a majority resolution identical to the species tree (Pamilo and Nei 1988, Rosenberg 2002, Degnan and Rosenberg 2009). The other two alternative gene tree resolutions will be equiprobable to one another and proportionally less than the majority resolution (Pamilo and Nei 1988, Rosenberg 2002, Degnan and Rosenberg 2009). To evaluate whether incongruence among our analyses was due to incomplete lineage sorting, we tested our 50p total evidence optimal gene trees for asymmetry among minority gene trees using an exact two-sided binomial test (Zwickl et al. 2014, Richart et al. 2016, Wang et al. 2017, Forthman et al. 2019). Gene trees were pruned to include a representative of three clades at the polytomy with the highest UCE recovery (*Galaesus hasticornis, Odontorhopala callosa, Anasa tristis*), as well as an outgroup (*Halyomorpha halys*).

## Results

### Data summary

For this study, we recovered 3,750–61,308 contigs across samples (mean = 13,284), with a mean length of 451 bp (Table S2). We recovered 9–55% of the targeted UCE loci (range across samples: 242–1,470 loci; mean = 1,040), with a mean length of 668 bp. We found that the TE-touchdown protocol recovered more contigs and loci on average (per sample: 4,922–61,308 contigs [mean = 18,017]; per sample: 242–1,470 loci [mean = 1,215]) than the TE protocol (per sample: 3,750–18,575 contigs [mean = 8,522]; per sample: 556–1,272 loci [mean = 858]) (Fig. 2). One sample produced an exceptional number of contigs compared to all others, while another produced very few UCE loci. Removal of these samples still produced qualitatively similar results (data not shown).

**Figure 2.**
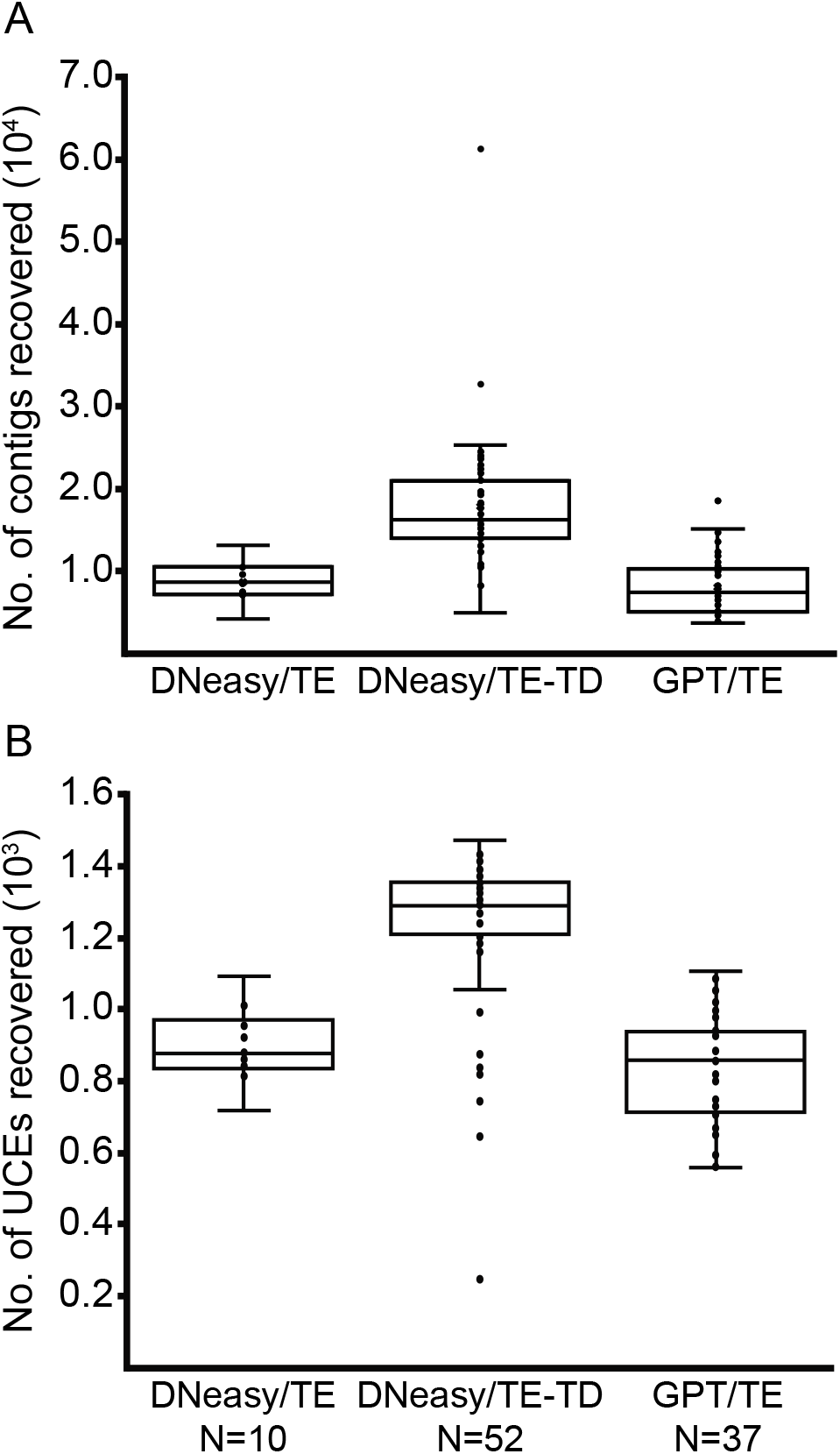
Contig (a) and UCE locus (b) recovery between different DNA extraction and target enrichment protocols. Abbreviations: DNeasy, Qiagen DNeasy Blood and Tissue kit (including QIAquick PCR Purification kit); GPT, Gentra Puregene Tissue kit; TE, (Forthman et al. n.d.) target enrichment protocol; TE-TD, TE-touchdown protocol.

Of the 16 dried museum samples in which genomic DNA was extracted, subjected to the TE-touchdown protocol, and sequenced, only five yielded >200 UCE loci. The remaining samples yielded too few or no UCE loci. Four of the successful samples were recently collected (2015–2016), while the fifth sample was much older (collected in 1946). For the more recent samples, contig recovery did not appear to be dependent on DNA extraction method, but UCE recovery did differ with the DNQIA extraction protocol yielding substantially more UCEs (Table S3). Similarly, contig length was not affected by the extraction protocol used, but extraction method had a noticeable effect on UCE locus length (Table S3). The older sample did not produce the lowest number of contigs (4,922; 9^th^ lowest), but it yielded the lowest number of UCE loci (242) in this study. Furthermore, this historical sample had smaller contig and UCE locus lengths on average (Table S3).

A summary of parsimony-informative sites and number of loci in each dataset are provided in Table S4. The most parsimony-informative UCE locus contained 1,079 informative sites, with the least informative locus having 16 informative sites. As expected, when datasets were constructed with higher locus informativeness thresholds, there was an increase in the proportion of parsimony-informative sites. There was also a decrease in the proportion of invariant sites, while the proportion of parsimony-uninformative sites were similar across all datasets.

After pruning outgroup taxa from our trees, symmetric distances were 0–8 among optimal summary coalescent trees, 0–8 among ML trees, and 2–12 when distances were calculated between coalescent and ML trees (Table S5). We recovered similar values when using trees with poorly supported branches (BS < 50%) collapsed (Table S6).

### Higher-level relationships of the Coreinae + Meropachyinae

Most reconstructed relationships within the Coreinae (including Meropachyinae) were congruent across all analyses (Figs. 3, S1–S8) and highly supported (Fig. 4). Some relationships were also consistently recovered across our estimates despite lower support (e.g., *Coreus* + *Cletus*; Cloresmini + Colpurini + Mictini) (Fig. 4). Across all analyses, Coreinae and Meropachyinae were not supported as monophyletic subfamilies. The meropachyine tribes Spathophorini and Merocorini were consistently recovered within coreine clades comprised of Nematopodini + Discogastrini and Acanthocerini + Chariesterini + Hypselonotini (part), respectively (Figs. 3, S1–S8), with moderate to high support (Fig. 4).

**Figure 3.**
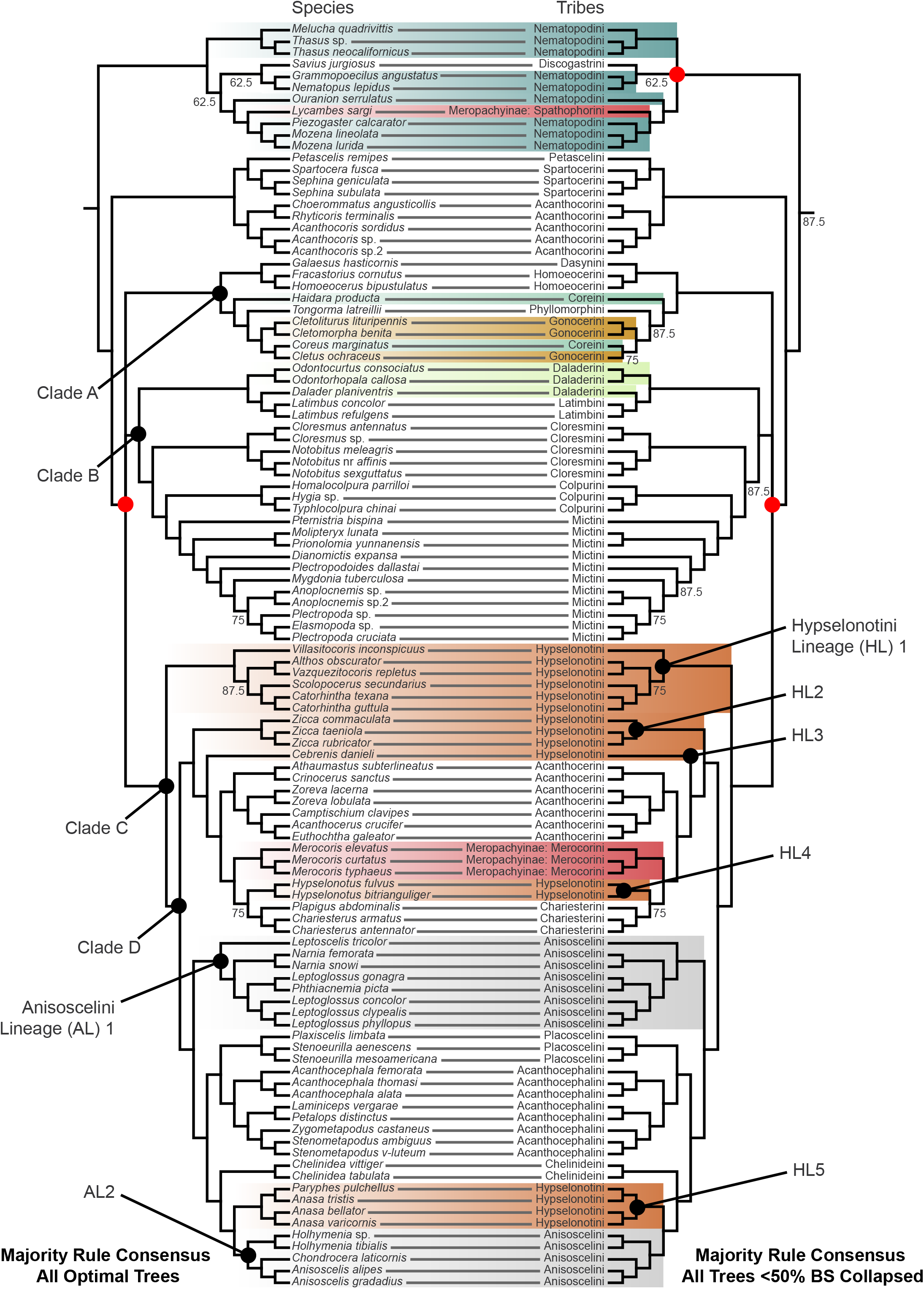
Majority rule consensus tree of all optimal trees (left) and all trees when branches with bootstrap support <50% are collapsed (right) across maximum likelihood and summary coalescent analyses (outgroups pruned for visualization) and datasets (i.e., taxon-sampling and parsimony informativeness). Species names are provided on the left tree, while the names of the corresponding tribes are provided on the right tree. Select tribes (including the subfamily Meropachyinae) that are non-monophyletic are color-coded. Red circles at nodes indicate the location of polytomies in the majority rule consensus trees. Numbers below branches indicate the proportion of all eight trees that recovered the corresponding clade; branches without numbers were recovered in 100% of trees.

**Figure 4.**
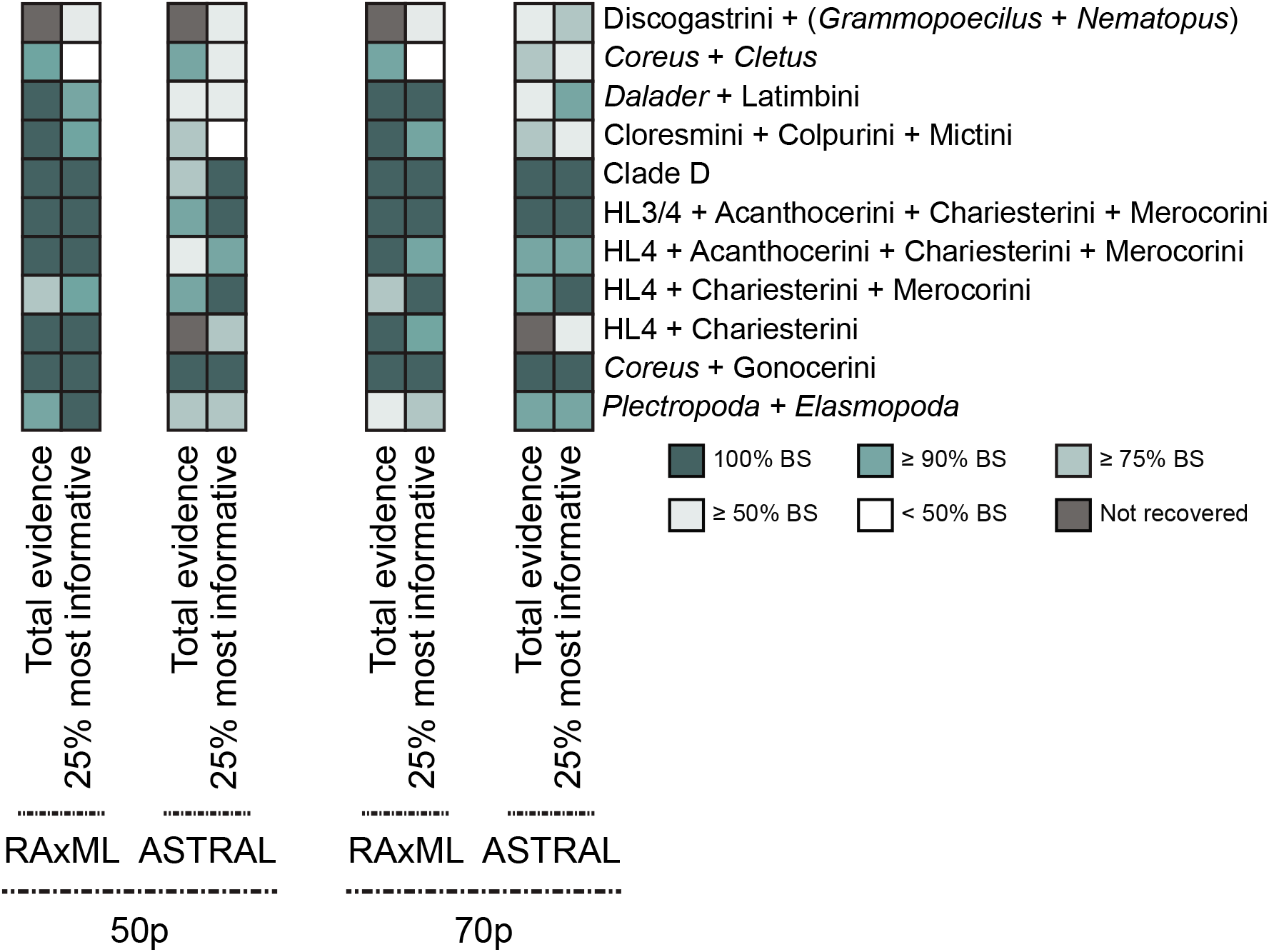
Bootstrap support for select Coreinae + Meropachyinae branches from Fig. 3 that exhibit conflict and/or low support across all analyses. Abbreviations: Bootstrap support, BS; Hypselonotini Lineage (HL).

The majority rule consensus tree of all estimated trees resulted in a single polytomy near the base of Coreinae + Meropachyinae (Fig. 3), which was driven by uncertainty in the phylogenetic placement of Clade A. This clade was either recovered as the sister to all members of Clade B in summary coalescent analyses (BS 97–100%) (Figs. S1–S4), sister to Clade B + Clade C in most ML analyses (BS 54–83%) (Figs. S5, S6, S8), or sister to Clade C in one ML analysis (BS = 83%) (Fig. S7). Given this incongruence among analyses, we tested if our estimated gene trees were consistent with the multispecies coalescent model. While a majority of our gene trees matched the species tree (46.64%), we detected asymmetry among our minority gene trees (33.40% and 18.26%; p < 0.05), suggesting incongruence was not driven by incomplete lineage sorting.

When poorly supported branches (BS < 50%) were collapsed in our estimated trees, we recovered an additional polytomy in our majority rule consensus tree among the Discogastrini + Nematopodini + Spathophorini (Fig. 3). The only sampled species of Discogastrini was more often recovered as the sister to the nematopodine genera *Nematopus* + *Grammopoecilius* with weak to moderate support (BS 55–76%) (Figs. S2–S4, S6, S8), but three analyses recovered this tribe as sister to all Nematopodini + Spathophorini with high support (BS = 100%) (Figs. S1, S5, S7). Furthermore, the position of the nematopodine genera *Melucha* + *Thasus* also varied across analyses, either recovered as the sister group of all other Nematopodini + Discogastrini + Spathophorini (BS = 100%) (Figs. S2–S4, S8), all remaining Nematopodini + Spathophorini (BS 97–100%) (Figs. S1, S6), or *Ouranion* + *Piezogaster* + *Mozena* + Spathophorini (BS 94–100%) (Figs. S5, S7).

The position of the meropachyine tribe Merocorini was typically found to be sister to *Hypselonotus* + Chariesterini with weak to high support (Figs. S2, S4–S8). In summary coalescent analyses that used all loci, Merocorini was recovered as the sister to *Hypselonotus* (Figs. S1, S3). However, support for this relationship exhibited drastically different values: support was 82% for our 50p total evidence dataset whereas it was 27% for the 70p total evidence dataset.

### Non-monophyly of coreine tribes

At the tribal level, we found support for the non-monophyly of several Coreinae tribes (Figs. 3, 4, S1–S8). Nematopodini was consistently not monophyletic with respect to Spathophorini or Spathophorini + Discogastrini. The Coreini were polyphelytic, with *Haidara* highly supported as the sister to Phyllomorphini + Gonocerini + *Coreus* and *Coreus* sister to *Cletus* with poor to high support. *Coreus* rendered Gonocerini paraphyletic with strong support. Weak to high support for a paraphyletic Daladerini with respect to Latimbini was observed across all analyses. Our results also recovered a polyphyletic Hypselonotini with five distinct lineages throughout Clade C, as well as a polyphyletic Anisoscelini with two lineages.

### Genus-level phylogenetic results

At shallower depths, our results supported most sampled genera as clades comprised of their respective conspecifics with strong support (Figs. 3, 4, S1–S8). Only a few genera were not monophyletic. *Leptoglossus* was paraphyletic with respect to *Phthiacnemia*, and *Paryphes* rendered *Anasa* as paraphyletic, both at high support. We also found evidence for a paraphyletic *Plectropoda* with respect to *Elasmopoda* with weak to high support.

## Discussion

We robustly reconstructed many shallow-level hemipteran relationships in the speciose coreid subfamily Coreinae. Our study recovered many highly support clades among and within coreine tribes and genera that were generally congruent across our maximum likelihood and summary coalescent analyses, although uncertainty still exists among the clades at deeper branches (i.e., clades A, B, and C in Fig. 3). The majority of our recovered relationships were novel, though a few were congruent with previous studies (e.g., Schaefer 1965, 1968, O’Shea 1980a, 1980b, Li 1996, 1997, Kieran et al. 2019, Forthman et al. 2019). Additionally, several tribes and genera — as well as the subfamilies Coreinae and Meropachyinae — were recovered as para-or polyphyletic, often with robust support; the taxonomic status of these groups should be evaluated further to revise classification.

### Evaluation of DNA extraction and target enrichment protocols

We observed a ~1.5– 2x increase in contig and UCE recovery when implementing our TE-touchdown approach. Such an increase may be due to the moderate reduction in hybridization temperature (i.e., from 65°C to 62°C). However, our TE-touchdown approach may have been confounded by early termination of our initial hybridization, although baits from the first hybridization should have been ineffective during the second attempt. Regardless, our improvement in locus recovery is consistent with other studies that have implemented a touchdown approach (e.g., Li et al. 2013, Paijmans et al. 2016), though not as high as has been reported in some studies (up to 8x increase; Li et al. 2013). This is likely due to our protocols conservatively reducing hybridization and washing temperatures (i.e., 62°C rather than 50°C). Further reductions in hybridization temperature during target capture may result in greater locus recovery with this and other invertebrate bait sets.

Our study suggests that recently collected, dried specimen material generally performs better than older samples in UCE target capture, consistent with other invertebrate UCE studies (e.g., Blaimer et al. 2016). However, based on our results, the choice of DNA extraction protocol may be important for locus recovery in dried material, including those that are more recently collected (>5 years old). While neither extraction protocol resulted in large differences in contig recovery or length, we found ~2x increase in the number of UCE loci recovered, as well as generally longer UCEs, when using the DNQIA protocol (Knyshov et al. 2019). Thus, for these types of samples, a protocol tailored for the extraction of smaller DNA fragments may improve overall yield in UCE studies, although the success of any extraction approaches with historical invertebrate samples may be variable (e.g., Blaimer et al. 2016). Although there is a general negative effect of sample age on target capture yields (Blaimer et al. 2016, McCormack et al. 2016), it is unclear what factors may have contributed to our limited success with our older samples; all dried samples were subjected to the same molecular protocols and conditions as the successfully sequenced sample, and equal amounts of starting tissue were used for genomic DNA extraction. It is possible that factors such as the rate of desiccation and/or preservation methods prior to curation may affect target capture results (Blaimer et al. 2016).

### Coreinae paraphyly and Meropachyinae polyphyly

Although Meropachyinae have rarely been included in phylogenetic analyses that sample Coreidae, there has been morphological and molecular evidence for the paraphyly of Coreinae with respect to this subfamily (Li 1996, 1997, Kieran et al. 2019, Forthman et al. 2019), which we corroborate. We found paraphyly of Nematopodini with respect to the meropachyine tribe Spathophorini, as in previous UCE studies (Kieran et al. 2019, Forthman et al. 2019). Additionally, the close relationship between Merocorini (Meropachyinae), Chariesterini, and *Hypselonotus* we recovered is largely consistent with previous cladistic (Li 1996, 1997) and non-cladistic (Schaefer 1965, Hepburn and Yonke 1971) studies. Both subfamilies have historically been delimited from the other coreid subfamilies by the presence of a dorsally sulcate tibia (see Forthman et al. 2019). In the taxonomic literature, the two subfamilies have been diagnosed from each other primarily by the presence (Meropachyinae) or absence (Coreinae) of an apical spine or tooth on the hind tibiae, as well as the shape of the hind femora and location of the metathoracic scent gland orifices. Our results indicate that these traits are likely homoplastic.

### Uncertain phylogenetic placement of clades A, B, and C

Our study finds robust support for a clade comprised of Dasynini, Homoeocerini, Coreini, Phyllomorphini, and Gonocerini (Clade A). The phylogenetic position of Clade A, however, remains uncertain. In our summary coalescent analyses, this clade was sister to Clade B (Daladerini, Latimbini, Cloresmini, Colpurini, and Mictini; see Fig. 3) with high support, while the ML analyses recovered two alternative topologies involving the large Clade C (Hypselonotini, Acanthocerini, Merocorini, Chariesterini, Anisoscelini, Placoscelini, Acanthocephalini, Chelinideini) with weaker support.

The internal branches around the polytomy we recovered are very short relative to other branches at deep nodes (Figs. S5–S8). The short, successive branches suggest that this region of the tree might be in an anomaly zone, i.e., a region of the species tree where discordant gene trees are more common than gene trees that are concordant with the species tree due to incomplete lineage sorting (Degnan and Rosenberg 2006, Liu and Edwards 2009). However, our test of minority gene tree asymmetry suggests that our estimated gene trees are inconsistent with the multispecies coalescent model. Thus, discordance around these branches is likely due to other processes.

### Non-monophyly of Nematopodini

A paraphyletic Nematopodini with respect to Spathophorini has also been supported by Kieran et al. (2019) and Forthman et al. (2019). However, in some of our analyses, Discogastrini was recovered within Nematopodini rather than as the sister group of Nematopodini + Spathophorini. This was not dependent on the analytical method or type of dataset used. Thus, Discogastrini may render Nematopodini (including Spathophorini) not monophyletic. To our knowledge, there are no previous hypotheses for a relationship between these three taxa.

Amyot and Serville (1843) included some members of Discogastrini within the Nematopodini based on the presence of enlarged, armed hind femora in males (although, in type images of several genera, the legs of Discogastrini appear slender and unarmed). Discogastrini was subsequently treated as a distinct group from the Nematopodini by Stål (1867), primarily due to the position of the abdominal spiracles. In his comparative morphological study, Schaefer (1965) included the Discogastrini, Homoeocerini, and Latimbini in his *Homoeocerus*-group (each tribe treated as a separate subgroup) based on the structure of the conjunctiva, metathoracic scent gland opening, laterotergites, and external genitalia (Nematopodini not examined). Our results are more in line with Amyot and Serville’s (1843) classification, although the shape and armature of the hind femora may not be synapomorphies for Discogastrini + Nematopodini (including Spathophorini).

### Clade A and the non-monophyly of Coreini and Gonocerini

The taxonomic history of the Coreini has undergone drastic changes over the last decade. Many of the genera once classified in Coreini are now treated as members of Hypselonotini and other tribes, which we followed here (see CoreoideaSF Team 2019). Our results provide robust support for the exclusion of these genera from the Coreini, but we do not find evidence for a monophyletic Coreini. The relatively close relationship of Coreini and Gonocerini — which is paraphyletic in our study — is congruent with Pan et al. (2007) and Pan et al.’s (2008) Cytb phylogenies but contradicts Li’s (1997) morphological and Fang and Nie’s (2007) COII phylogenies. Schaefer (1965) placed the Coreini (which included genera now in other tribes) with several others (including Gonocerini and Dasynini) in his large *Coreus*-group. Our phylogenetic result provides limited support to Schaefer’s (1965) study but does not recognize the placement of many other tribes within his *Coreus*-group. Thus, the characters Schaefer (1965) used to diagnose and describe his *Coreus*-group and subgroups are likely plesiomorphic or homoplastic synapomorphies based on our molecular phylogenetic hypothesis.

The tribe Phyllomorphini has not been included in phylogenetic analyses. Past pre-cladistic morphological studies led some authors to propose Phyllomorphini as a distinct subfamily due to the absence of a dorsal tibial sulcus, as well as several genitalic and abdominal traits (Schaefer 1965, Ahmad 1970, 1979). However, we find support for the inclusion of this tribe within the Coreidae.

Lastly, the sister group relationship between Dasynini and Homoeocerini recovered from our analysis is novel. Schaefer (1965) did not consider these tribes to belong to the same group in his morphological study. Li (1997) included several species of Dasynini and found this tribe to be paraphyletic, but none of the sampled species were found to be closely related to Homoeocerini.

### Novel relationships of Clade B, the paraphyly of Daladerini, and phylogenetic position of Colpurini

The relationships recovered within our well-supported Clade B do not strictly agree with previous studies (Schaefer 1965, Li 1997, Fang and Nie 2007). Daladerini and Latimbini were recovered as the sister of all other tribes within Clade B. We found support for a clade comprised of Cloresmini, as well as its sister group relationship with Colpurini + Mictini that has not been previously proposed. Schaefer (1965) included Cloresmini within his *Coreus*-subgroup C with other tribes not recovered in this clade (although Mictini was included in a separate *Coreus*-subgroup within a larger *Coreus*-group).

Amyot and Serville (1843) classified *Dalader* in the same family-group as genera from Mictini, but it was subsequently treated as a distinct group by Stål (1873). Li’s (1997) morphological phylogenetic hypothesis suggests that the Daladerini are sister to Acanthocerini and Acanthocephalini, in support of Stål’s (1873) treatment of the tribe as separate from the Mictini. Our results also support the exclusion of the sampled daladerine genera from Mictini but find new evidence for the paraphyly of this tribe with respect to Latimbini. Schaefer (1965) assigned Daladerini to his *Coreus*-subgroup B; he assigned the Latimbini to the *Homoeocerus*-group but recognized that the position of Latimbini was uncertain.

Our results for the phylogenetic placement of Colpurini and treatment as a tribe within Coreinae is contradictory with all other studies (Štys 1964, Kumar 1965, Schaefer 1965, Li 1997, Ahmad 1970). Past studies have characterized the Colpurini as “primitive” but with many characters (primarily genitalia) suggesting an “intermediate” phylogenetic position between Pseudophloeinae, Hydarinae, and other Coreinae (see Štys 1964, Kumar 1965, Schaefer 1965, Li 1996, 1997, Ahmad 1970). Thus, our results are novel and suggest that genitalic features, as well as external features, should be re-evaluated in light of our molecular hypothesis.

### Polyphyly of Hypselonotini

With 356 species, the Hypselonotini is the most speciose tribe within the family Coreidae. This tribe has not been formally described or diagnosed even when it was first recognized by Bergroth (1913). Over the last three decades, a number of genera currently recognized within the Hypselonotini (CoreoideaSF Team 2019) have been previously treated as members of the Coreini (e.g., Brailovsky 1988, 1990, 1995, 2016, Packauskas 1994). Members of this tribe have not been included in published phylogenetic analyses and, to our knowledge, appear to have only been examined by Schaefer (1965) in his comparative morphological study (treated as part of Coreini). Our study supports the exclusion of the sampled hypselonotine genera from the Coreini but not the monophyly of this large tribe; five independent lineages were robustly supported in our analyses. Of these, only the taxonomic position of *Hypselonotus* within the *Hypselonotus* + Chariesterini + Merocorini was not congruent across analyses. It is evident from our results that the taxonomic status of Hypselonotini is in further need of evaluation. Including additional genera of this tribe will provide further insights into the extent of hypselonotine polyphyly.

### Polyphyly of Anisoscelini

The tribe Anisoscelini is a moderately-sized group, with 183 species that exhibit a diversity of color patterns and morphology, particularly with the shape and size of the foliaceous expansions on the hind tibiae. Members of Anisoscelini were once divided among two tribes (Anisosceledini [or Anisoscelini] and Leptoscelidini), primarily based on the presence or absence of hind tibial expansions (e.g., Stål 1867, Schaefer 1965, Packauskas 1994). Schaefer (1965, 1968) classified both of these former tribes as members of the *Acanthocephala*-group with Acanthocephalini and Placoscelini. We find some support for Schaefer’s (1965, 1968) scheme, but we do not support his exclusion of Chelinideini and Hypselonotini from the *Acanthocephala*-group. The two distinct anisosceline lineages we recovered do not appear to correspond to previously proposed tribal classifications and phylogenetic hypotheses nor appear to be separated based on the presence of tibial expansions. Like Hypselonotini, careful evaluation of this tribe is needed to understand the extent of anisosceline polyphyly.

### Phylogenetic position of Chelinideini

We support a sister group relationship between Chelinideini and our Hypselonotini Lineage 5 + Anisoscelini Lineage 2 clade (Fig. 3), contrary to previous studies. Li’s (1997) phylogenetic hypothesis found a close relationship between Chelinideini and Homoeocerini. Based on a survey of genitalic morphology, Schaefer (1965) placed this tribe in a *Coreus* subgroup (subgroup C) that included Gonocerini, Acanthocorini (part), Cloresmini, and Coreini. In fact, Chelinideini was once classified within the Coreini, but was elevated to tribal rank by Blatchley (1926). Thus, our results on the phylogenetic position of Chelinideini are novel and require further comparative work to identify and test potential synapomorphies.

### Paraphyletic genera

The genera *Plectropoda* and *Elasmopoda* are two of several genera comprising the *Elasmopoda* complex (Linnavuori 1978, O’Shea 1980c) and share many morphological similarities. Both genera have been treated as separate groups since Stål (1873), but some species have historically experienced changes in generic assignment between these two groups. Linnavuori (1978) revised the *Elasmopoda* complex but noted that some features used to diagnosis *Elasmopoda* are also observed in some *Plectropoda* species. This may suggest that one or both genera are not monophyletic, and our study supports the paraphyly of *Plectropoda* with respect to *Elasmopoda*. This indicate that the taxonomic limits of these genera should be evaluated further.

The genus *Leptoglossus* is an agriculturally important group (Schaefer and Mitchell 1983, Jankevicius et al. 1993, Fernandes et al. 2015), with many of the 62 species distributed throughout the New World, while two species, *L. gonagra* and *L. occidentalis*, occur worldwide (CoreoideaSF Team 2019). Our study recovered a paraphyletic *Leptoglossus*, with *L. gonagra* recovered as sister to *Phthiacnemia picta*, both of which were sampled from the New World. Based on our survey of the taxonomic literature and the Coreoidea Species File (CoreoideaSF Team 2019), these two genera have never been treated as or hypothesized to be the same. Allen (1969) created two divisions within *Leptoglossus* — Divisions A and B. While we do not support the monophyly of this genus, we do find limited support for Allen’s (1969) separate treatment of his Division A (i.e., *L. gonagra*) from those of Division B (i.e., *L. phyllopus, L. clypealis, L. concolor*).

The hypselonotine genus *Anasa* is a large genus comprised of 77 New World species (CoreoideaSF Team 2019), with several of economic importance (Schaefer and Mitchell 1983, Fernandes et al. 2015). We also found evidence for the paraphyly of *Anasa* with respect to the hypselonotine genus *Paryphes*. The close relationship between these two genera have not been previously tested. Stål (1867) provided a framework to separate the two genera based on the curvature of the head and the structure of the antennal segments. In this same publication, Stål (1867) transferred a single species from *Paryphes* to *Anasa*. Since then, the taxonomic literature has recognized these two genera as distinct without suggestion that they are potentially the same.

## Conclusion

The results of our phylogenomic analysis suggest para-and polyphyly of several genera, tribes, and subfamilies of Coreidae, indicating that the taxonomic classification of this diverse family and its largest subfamily, the Coreinae, is in critical need of evaluation and future revision. We were able to robustly reconstruct relationships at shallow phylogenetic scales within the coreid subfamily Coreinae, demonstrating that invertebrate UCEs are suitable at a variety of scales. Additionally, our results suggest that DNA extraction protocols designed to capture shorter, degraded DNA fragments in dried museum material and lower target capture hybridization temperatures may increase the number and length of UCE loci in this and potentially other invertebrate bait sets.

## Acknowledgements

We thank the village it took to make this study happen. We thank the following individuals and institutions that contributed specimens used in this study: Christiane Weirauch, Dimitri Forero, Ummat Somjee, Takahisa Miyatake, Jason Cryan, Mark Deyrup, Wei Song Hwang, Li You, Oliver Keller, John Leavengood, Sam Noble Oklahoma Museum of Natural History, Field Museum of Natural History, California State Collection of Arthropods, and National Museums of Kenya. Harry Brailovsky assisted with the identification of a few coreid species. Habitus images of vouchers were prepared by Cassie Bakus, Nathan Friedman, Courtney Gormley, Emma Matzinger, and Caroline Miller. Nathan Friedman and Caroline Miller assisted with molecular protocols. We thank Travis Glenn and Troy Kieran (University of Georgia) for intensive training in library preparation and target enrichment. Gavin Naylor and Shannon Corrigan provided training on Covaris ultrasonication. Collection of Florida (USA) specimens was supported by Archbold Biological Station, Kai Kai Farms, Rotary Community Garden and Food Forest of Coral Springs, and the Florida Park Service and Division of Recreation and Parks (permits #05051610 and #09011720). Bob McCleery (University of Florida), Cebisile N. Magagula (University of Swaziland), and the Savannah Research Center facilitated collection of specimens in the Kingdom of eSwatini (formerly known as Swaziland). Ezemvelo KZN Wildlife (permit #OP 172/2018) and the iSimangaliso Wetland Park Authority (World Heritage Site) facilitated specimen collection in South Africa. All Out Africa provided additional assistance in conducting research in the Kingdom of eSwatini and South Africa. Some Thailand specimens were acquired under NSF DEB 0542864 (awarded to Michael Sharkey and Brian Brown). Collection of Singapore specimens (National Park permit #NP/RP17-012) was supported by the Lee Kong Chian Natural History Museum fellowship and the Nation Science Foundation grant OISE-1614015 (awarded to Zachary Emberts). Collection of Australian specimens (Regulation 17 permit # 01-000204-1) was supported by a University of Florida Research Abroad for Doctoral Students (awarded to Zachary Emberts). Additional samples collected were supported by funding awarded to Ummat Somjee: Society of the Study of Evolution Rosemary Grant Award; Smithsonian Tropical Research Institute Short Term Fellowship; and Systematics, Evolution, and Biodiversity Endowment Award (Entomological Society of America). This study was funded by the National Science Foundation IOS-1553100 (awarded to C.W. Miller).

## Data accessibility

Sequence read files are available on NCBI’s Sequence Read Archive under BioProject PRJNA546248.

