## Supporting tables and figures for "Phylogenomics of the leaf-footed bug subfamily Coreinae (Hemiptera: Coreidae): applicability of ultraconserved elements at shallower depths"

### Supporting Information

Table S1. Taxon sampling and summary information on molecular protocols. Abbreviations: 95–100% EtOH, ethanol; GPT, Gentra Puregene Tissue kit; DNeasy, Qiagen QIAquick PCR Purification kit; TE, Forthman et al. (accepted) target enrichment protocol; TE-TD, TE-touchdown protocol.

| Family | Subfamily | Tribe | Genus | Species | Storage | Collection year | Tissue sampled | DNA extraction | Sonication | Target capture protocol |
|---|---|---|---|---|---|---|---|---|---|---|
| Coreidae | Coreinae | Acanthocephalini | *Acanthocephala* | *alata* | EtOH | 2010 | Legs | GPT | Bioruptor | TE |
| Coreidae | Coreinae | Acanthocephalini | *Laminiceps* | *vergarae* | EtOH | 2011 | Abdomen | DNeasy | Covaris | TE-TD |
| Coreidae | Coreinae | Acanthocephalini | *Petalops* | *distinctus* | EtOH | 2011 | Legs | DNeasy | Covaris | TE-TD |
| Coreidae | Coreinae | Acanthocephalini | *Stenometapodus* | *ambiguus* | EtOH | 2010 | Legs | DNeasy | Covaris | TE-TD |
| Coreidae | Coreinae | Acanthocephalini | *Stenometapodus* | *v-luteum* | EtOH | 2010 | Legs | DNeasy | Covaris | TE-TD |
| Coreidae | Coreinae | Acanthocephalini | *Zygometapodus* | *castaneus* | EtOH | 2011 | Legs | DNeasy | Covaris | TE-TD |
| Coreidae | Coreinae | Acanthocerini | *Acanthocerus* | *crucifer* | EtOH | 2017 | Legs | DNeasy | Covaris | TE-TD |
| Coreidae | Coreinae | Acanthocerini | *Athaumastus* | *subterlineatus* | Dried | 2016 | Legs | DNeasy | Covaris | TE-TD |
| Coreidae | Coreinae | Acanthocerini | *Camptischium* | *clavipes* | Frozen | 2017 | Abdomen | DNeasy | Covaris | TE-TD |
| Coreidae | Coreinae | Acanthocerini | *Crinocerus* | *sanctus* | EtOH | 2010 | Legs & abdomen | DNeasy | Covaris | TE-TD |
| Coreidae | Coreinae | Acanthocerini | *Euthochtha* | *galeator* | EtOH | 2016 | Abdomen | DNeasy | Bioruptor | TE |
| Coreidae | Coreinae | Acanthocerini | *Zoreva* | *lacerna* | EtOH | 2010 | Legs & abdomen | DNeasy | Covaris | TE-TD |
| Coreidae | Coreinae | Acanthocerini | *Zoreva* | *lobulata* | Frozen | 2017 | Thorax, legs, & abdomen | DNeasy | Covaris | TE-TD |
| Coreidae | Coreinae | Acanthocorini | *Acanthocoris* | *sordidus* | EtOH | unknown | Whole body | GPT | Bioruptor | TE |
| Coreidae | Coreinae | Acanthocorini | *Acanthocoris* | sp. | Silica beads | 2015 | Thorax & legs | GPT | Bioruptor | TE |
| Coreidae | Coreinae | Acanthocorini | *Acanthocoris* | sp.2 | EtOH | 2008 | Legs | GPT | Bioruptor | TE |
| Coreidae | Coreinae | Acanthocorini | *Choerommatus* | *angusticollis* | EtOH | 2013 | Legs & abdomen | GPT | Bioruptor | TE |
| Coreidae | Coreinae | Acanthocorini | *Rhyticoris* | *terminalis* | EtOH | 2018 | Abdomen | DNeasy | Covaris | TE-TD |

| | | | | | | | | | | |
|---|---|---|---|---|---|---|---|---|---|---|
| Coreidae | Coreinae | Anisoscelini | *Anisoscelis* | *gradadius* | EtOH | 2013 | Legs & abdomen | GPT | Bioruptor | TE |
| Coreidae | Coreinae | Anisoscelini | *Chondrocera* | *laticornis* | EtOH | 2018 | Abdomen | DNeasy | Covaris | TE-TD |
| Coreidae | Coreinae | Anisoscelini | *Holhymenia* | sp. | EtOH | 2010 | Legs | GPT | Bioruptor | TE |
| Coreidae | Coreinae | Anisoscelini | *Holhymenia* | *tibialis* | EtOH | unknown | Abdomen | DNeasy | Covaris | TE-TD |
| Coreidae | Coreinae | Anisoscelini | *Leptoglossus* | *clypealis* | EtOH | 2016 | Legs & abdomen | DNeasy | Bioruptor | TE |
| Coreidae | Coreinae | Anisoscelini | *Leptoglossus* | *concolor* | EtOH | 2009 | Legs | GPT | Bioruptor | TE |
| Coreidae | Coreinae | Anisoscelini | *Leptoglossus* | *gonagra* | EtOH | 2018 | Legs & abdomen | DNeasy | Covaris | TE-TD |
| Coreidae | Coreinae | Anisoscelini | *Leptoglossus* | *phyllopus* | EtOH | 2016 | Legs & abdomen | GPT | Bioruptor | TE |
| Coreidae | Coreinae | Anisoscelini | *Leptoscelis* | *tricolor* | EtOH | 2008 | Legs | GPT | Bioruptor | TE |
| Coreidae | Coreinae | Anisoscelini | *Narnia* | *femorata* | EtOH | 2009 | Legs | GPT | Bioruptor | TE |
| Coreidae | Coreinae | Anisoscelini | *Narnia* | *snowi* | EtOH | 2016 | Abdomen | DNeasy | Bioruptor | TE |
| Coreidae | Coreinae | Anisoscelini | *Phthiacnemia* | *picta* | EtOH | 2017 | Abdomen | DNeasy | Bioruptor | TE |
| Coreidae | Coreinae | Chariesterini | *Chariesterus* | *antennator* | EtOH | 2016 | Legs & abdomen | GPT | Bioruptor | TE |
| Coreidae | Coreinae | Chariesterini | *Chariesterus* | *armatus* | EtOH | 2010 | Thorax, legs, & abdomen | DNeasy | Covaris | TE-TD |
| Coreidae | Coreinae | Chariesterini | *Plapigus* | *abdominalis* | EtOH | 2013 | Legs & abdomen | GPT | Bioruptor | TE |
| Coreidae | Coreinae | Chelinideini | *Chelinidea* | *tabulata* | EtOH | 2016 | Abdomen | DNeasy | Bioruptor | TE |
| Coreidae | Coreinae | Chelinideini | *Chelinidea* | *vittiger* | EtOH | 2016 | Abdomen | DNeasy | Bioruptor | TE |
| Coreidae | Coreinae | Cloresmini | *Cloresmus* | *antennatus* | EtOH | 2017 | Legs & abdomen | DNeasy | Covaris | TE-TD |
| Coreidae | Coreinae | Cloresmini | *Cloresmus* | sp. | EtOH | 2009 | Legs & abdomen | DNeasy | Covaris | TE-TD |
| Coreidae | Coreinae | Cloresmini | *Notobitus* | *meleagris* | Dried | 2015 | Legs | DNeasy | Covaris | TE-TD |
| Coreidae | Coreinae | Cloresmini | *Notobitus* | nr *affinis* | Dried | 2015 | Legs & abdomen | DNQIA | Covaris | TE-TD |
| Coreidae | Coreinae | Cloresmini | *Notobitus* | *sexguttatus* | EtOH | 2007 | Abdomen | DNeasy | Covaris | TE-TD |
| Coreidae | Coreinae | Colpurini | *Homalocolpura* | *parrilloi* | Dried | 1946 | Legs & abdomen | DNeasy | Unsheared | TE-TD |
| Coreidae | Coreinae | Colpurini | *Hygia* | sp. | EtOH | 2006 | Head & pygophore | DNeasy | Covaris | TE-TD |
| Coreidae | Coreinae | Colpurini | *Typhlocolpura* | *chinai* | EtOH | 2006 | Legs & abdomen | GPT | Bioruptor | TE |
| Coreidae | Coreinae | Coreini | *Coreus* | *marginatus* | EtOH | 2017 | Legs & abdomen | DNeasy | Covaris | TE-TD |
| Coreidae | Coreinae | Coreini | *Haidara* | *producta* | EtOH | 2005 | Whole body | DNeasy | Covaris | TE-TD |
| Coreidae | Coreinae | Daladerini | *Dalader* | *planiventris* | EtOH | 2006 | Legs | DNeasy | Covaris | TE-TD |

| | | | | | | | | | | |
|---|---|---|---|---|---|---|---|---|---|---|
| Coreidae | Coreinae | Daladerini | *Odontocurtus* | *consociatus* | EtOH | 2008 | Abdomen | DNeasy | Covaris | TE-TD |
| Coreidae | Coreinae | Daladerini | *Odontorhopala* | *callosa* | EtOH | 2008 | Abdomen | DNeasy | Covaris | TE-TD |
| Coreidae | Coreinae | Dasynini | *Galaesus* | *hasticornis* | EtOH | 2018 | Legs & abdomen | DNeasy | Covaris | TE-TD |
| Coreidae | Coreinae | Discogastrini | *Savius* | *jurgiosus* | EtOH | 2008 | Legs & abdomen | GPT | Bioruptor | TE |
| Coreidae | Coreinae | Gonocerini | *Cletoliturus* | *lituripennis* | EtOH | 2018 | Whole body | DNeasy | Covaris | TE-TD |
| Coreidae | Coreinae | Gonocerini | *Cletomorpha* | *benita* | EtOH | 2006 | Whole body | DNeasy | Covaris | TE-TD |
| Coreidae | Coreinae | Gonocerini | *Cletus* | *ochraceus* | EtOH | 2008 | Whole body | GPT | Bioruptor | TE |
| Coreidae | Coreinae | Homoeocerini | *Fracastorius* | *cornutus* | Dried | 2015 | Legs & abdomen | DNQIA | Covaris | TE-TD |
| Coreidae | Coreinae | Homoeocerini | *Homoeocerus* | *bipustulatus* | EtOH | 2008 | Abdomen | GPT | Bioruptor | TE |
| Coreidae | Coreinae | Hypselonotini | *Althos* | *obscurator* | EtOH | 2010 | Head, thorax, & legs | DNeasy | Covaris | TE-TD |
| Coreidae | Coreinae | Hypselonotini | *Anasa* | *bellator* | EtOH | 2010 | Legs & abdomen | GPT | Bioruptor | TE |
| Coreidae | Coreinae | Hypselonotini | *Anasa* | *tristis* | EtOH | 2017 | Abdomen | DNeasy | Covaris | TE-TD |
| Coreidae | Coreinae | Hypselonotini | *Anasa* | *varicornis* | EtOH | 2013 | Legs & abdomen | GPT | Bioruptor | TE |
| Coreidae | Coreinae | Hypselonotini | *Catorhintha* | *guttula* | EtOH | 2016 | Whole body | GPT | Bioruptor | TE |
| Coreidae | Coreinae | Hypselonotini | *Catorhintha* | *texana* | EtOH | 2015 | Legs & abdomen | GPT | Bioruptor | TE |
| Coreidae | Coreinae | Hypselonotini | *Cebrenis* | *danieli* | EtOH | 2013 | Whole body | GPT | Bioruptor | TE |
| Coreidae | Coreinae | Hypselonotini | *Hypselonotus* | *bitrianguliger* | EtOH | 2014 | Legs & abdomen | GPT | Bioruptor | TE |
| Coreidae | Coreinae | Hypselonotini | *Hypselonotus* | *fulvus* | EtOH | 2014 | Legs & abdomen | GPT | Bioruptor | TE |
| Coreidae | Coreinae | Hypselonotini | *Paryphes* | *pulchellus* | EtOH | 2010 | Thorax & legs | DNeasy | Covaris | TE-TD |
| Coreidae | Coreinae | Hypselonotini | *Scolopocerus* | *secundarius* | EtOH | 2017 | Whole body | DNeasy | Covaris | TE-TD |
| Coreidae | Coreinae | Hypselonotini | *Vazquezitocoris* | *repletus* | EtOH | 2017 | Whole body | DNeasy | Covaris | TE-TD |
| Coreidae | Coreinae | Hypselonotini | *Villasitocoris* | *inconspicuus* | EtOH | 2017 | Abdomen | DNeasy | Covaris | TE-TD |
| Coreidae | Coreinae | Hypselonotini | *Zicca* | *commaculata* | EtOH | 2008 | Legs & abdomen | GPT | Bioruptor | TE |
| Coreidae | Coreinae | Hypselonotini | *Zicca* | *rubricator* | EtOH | 2008 | Whole body | GPT | Bioruptor | TE |
| Coreidae | Coreinae | Hypselonotini | *Zicca* | *taeniola* | EtOH | 2017 | Whole body | GPT | Bioruptor | TE |
| Coreidae | Coreinae | Latimbini | *Latimbus* | *concolor* | EtOH | 2018 | Thorax & abdomen | DNeasy | Covaris | TE-TD |
| Coreidae | Coreinae | Latimbini | *Latimbus* | *refulgens* | EtOH | 2014 | Legs & abdomen | DNeasy | Covaris | TE-TD |
| Coreidae | Coreinae | Mictini | *Anoplocnemis* | sp.2 | EtOH | 2008 | Legs | GPT | Bioruptor | TE |

| | | | | | | | | | | |
|---|---|---|---|---|---|---|---|---|---|---|
| Coreidae | Coreinae | Mictini | *Dianomictis* | *expansa* | EtOH | 2008 | Legs | DNeasy | Covaris | TE-TD |
| Coreidae | Coreinae | Mictini | *Elasmopoda* | sp. | Silica beads | 2015 | Legs | GPT | Bioruptor | TE |
| Coreidae | Coreinae | Mictini | *Molipteryx* | *lunata* | EtOH | 2012 | Legs | DNeasy | Covaris | TE-TD |
| Coreidae | Coreinae | Mictini | *Plectropoda* | *cruciata* | EtOH | 2013 | Legs | DNeasy | Covaris | TE-TD |
| Coreidae | Coreinae | Mictini | *Plectropoda* | sp. | EtOH | 2013 | Legs | GPT | Bioruptor | TE |
| Coreidae | Coreinae | Mictini | *Plectropodoides* | *dallastai* | EtOH | 2009 | Legs | DNeasy | Covaris | TE-TD |
| Coreidae | Coreinae | Mictini | *Prionolomia* | *yunnanensis* | EtOH | 2017 | Legs | DNeasy | Covaris | TE-TD |
| Coreidae | Coreinae | Mictini | *Pternistria* | *bispina* | EtOH | 2011 | Legs | DNeasy | Covaris | TE-TD |
| Coreidae | Coreinae | Nematopodini | *Grammopoecilus* | *angustatus* | EtOH | 2010 | Legs | DNeasy | Covaris | TE-TD |
| Coreidae | Coreinae | Nematopodini | *Melucha* | *quadrivittis* | EtOH | 2013 | Legs | GPT | Bioruptor | TE |
| Coreidae | Coreinae | Nematopodini | *Mozena* | *lurida* | EtOH | 2010 | Legs | GPT | Bioruptor | TE |
| Coreidae | Coreinae | Nematopodini | *Nematopus* | *lepidus* | EtOH | 2013 | Legs | GPT | Bioruptor | TE |
| Coreidae | Coreinae | Nematopodini | *Ouranion* | *serrulatus* | frozen | 2017 | Legs | GPT | Bioruptor | TE |
| Coreidae | Coreinae | Nematopodini | *Piezogaster* | *calcarator* | EtOH | 2018 | Legs | DNeasy | Covaris | TE-TD |
| Coreidae | Coreinae | Nematopodini | *Thasus* | sp. | EtOH | 2013 | Legs | GPT | Bioruptor | TE |
| Coreidae | Coreinae | Petascelini | *Petascelis* | *remipes* | Silica beads | 2015 | Legs | GPT | Bioruptor | TE |
| Coreidae | Coreinae | Phyllomorphini | *Tongorma* | *latreilii* | EtOH | 2018 | Whole body | DNeasy | Covaris | TE-TD |
| Coreidae | Coreinae | Placoscelini | *Plaxiscelis* | *limbata* | EtOH | 2014 | Thorax & abdomen | GPT | Bioruptor | TE |
| Coreidae | Coreinae | Placoscelini | *Stenoeurilla* | *mesoamericana* | EtOH | 2013 | Abdomen | DNeasy | Bioruptor | TE |
| Coreidae | Coreinae | Spartocerini | *Sephina* | *geniculata* | EtOH | 2008 | Abdomen | DNeasy | Bioruptor | TE |
| Coreidae | Coreinae | Spartocerini | *Sephina* | *subulata* | EtOH | 2008 | Legs | DNeasy | Covaris | TE-TD |
| Coreidae | Coreinae | Spartocerini | *Spartocera* | *fusca* | EtOH | 2017 | Head, thorax, & legs | DNeasy | Bioruptor | TE |
| Coreidae | Meropachyinae | Merocorini | *Merocoris* | *curtatus* | EtOH | 2017 | Whole body | DNeasy | Covaris | TE-TD |
| Coreidae | Meropachyinae | Merocorini | *Merocoris* | *elevatus* | EtOH | 2010 | Whole body | DNeasy | Covaris | TE-TD |
| Coreidae | Meropachyinae | Merocorini | *Merocoris* | *typhaeus* | EtOH | 2017 | Whole body | DNeasy | Bioruptor | TE |

930

Table S2. Summary data for sequence reads, contigs, and ultraconserved element loci generated in this study.

| Family | Subfamily | Tribe | Genus | Species | Paired Reads | Reads passed QC | Contigs | Total bp | Mean contig length | Min contig length | Max contig length | UCE loci | % UCE recovered | Mean UCE length | Min UCE length | Max UCE length |
|---|---|---|---|---|---|---|---|---|---|---|---|---|---|---|---|---|
| Coreidae | Coreinae | Acanthocephalini | *Acanthocephala* | *alata* | 17030816 | 1761888 | 5144 | 2366942 | 460 | 194 | 2659 | 854 | 31.95% | 564 | 201 | 2659 |
| Coreidae | Coreinae | Acanthocephalini | *Laminiceps* | *vergarae* | 12909816 | 1721730 | 10466 | 4815312 | 460 | 198 | 2887 | 1272 | 47.59% | 634 | 202 | 2775 |
| Coreidae | Coreinae | Acanthocephalini | *Petalops* | *distinctus* | 15018864 | 2088428 | 12294 | 5777112 | 470 | 195 | 3171 | 1212 | 45.34% | 705 | 201 | 3171 |
| Coreidae | Coreinae | Acanthocephalini | *Stenometapodus* | *ambiguus* | 9421912 | 1274716 | 8267 | 4125586 | 499 | 194 | 3396 | 1200 | 44.89% | 672 | 203 | 2836 |
| Coreidae | Coreinae | Acanthocephalini | *Stenometapodus* | *v-luteum* | 23084240 | 2928640 | 16983 | 8117379 | 478 | 184 | 4175 | 1279 | 47.85% | 756 | 201 | 3032 |
| Coreidae | Coreinae | Acanthocephalini | *Zygometapodus* | *castaneus* | 18621536 | 2567350 | 14965 | 6702077 | 448 | 189 | 5896 | 1308 | 48.93% | 698 | 202 | 3011 |
| Coreidae | Coreinae | Acanthocerini | *Acanthocerus* | *crucifer* | 39593704 | 5312354 | 15340 | 5910208 | 385 | 182 | 2633 | 835 | 31.24% | 488 | 201 | 2633 |
| Coreidae | Coreinae | Acanthocerini | *Athaumastus* | *subterlineatus* | 57094592 | 5179960 | 17167 | 6838761 | 398 | 194 | 3143 | 645 | 24.13% | 433 | 203 | 1580 |
| Coreidae | Coreinae | Acanthocerini | *Camptischium* | *clavipes* | 76156480 | 10903458 | 19860 | 7464361 | 376 | 183 | 2798 | 743 | 27.80% | 470 | 201 | 2191 |
| Coreidae | Coreinae | Acanthocerini | *Crinocerus* | *sanctus* | 32775616 | 3608494 | 14742 | 6868069 | 466 | 199 | 3227 | 1283 | 48.00% | 730 | 201 | 2900 |
| Coreidae | Coreinae | Acanthocerini | *Euthochtha* | *galeator* | 21289056 | 2509972 | 8525 | 4371314 | 513 | 191 | 4086 | 814 | 30.45% | 829 | 201 | 3212 |
| Coreidae | Coreinae | Acanthocerini | *Zoreva* | *lacerna* | 25366184 | 3663304 | 21033 | 9605500 | 457 | 194 | 6279 | 1208 | 45.19% | 683 | 202 | 2733 |
| Coreidae | Coreinae | Acanthocerini | *Zoreva* | *lobulata* | 55386416 | 5290149 | 16250 | 6833729 | 421 | 187 | 3655 | 825 | 30.86% | 531 | 201 | 2166 |
| Coreidae | Coreinae | Acanthocorini | *Acanthocoris* | *sordidus* | 28851552 | 2702481 | 7458 | 3727512 | 500 | 195 | 2842 | 1018 | 38.08% | 678 | 202 | 2776 |
| Coreidae | Coreinae | Acanthocorini | *Acanthocoris* | sp. | 17775496 | 2257165 | 7698 | 3755927 | 488 | 201 | 3812 | 746 | 27.91% | 738 | 202 | 3047 |
| Coreidae | Coreinae | Acanthocorini | *Acanthocoris* | sp.2 | 26052120 | 2248712 | 4854 | 2198119 | 453 | 201 | 3439 | 660 | 24.69% | 590 | 201 | 3439 |
| Coreidae | Coreinae | Acanthocorini | *Choerommatus* | *angusticollis* | 22812224 | 2742939 | 11910 | 5838231 | 490 | 193 | 4724 | 993 | 37.15% | 872 | 202 | 3327 |
| Coreidae | Coreinae | Acanthocorini | *Rhyticoris* | *terminalis* | 14621664 | 1744098 | 15248 | 6483622 | 425 | 189 | 3971 | 1337 | 50.02% | 656 | 201 | 3971 |
| Coreidae | Coreinae | Anisoscelini | *Anisoscelis* | *gradadius* | 29365416 | 4125411 | 13621 | 6248865 | 459 | 194 | 3354 | 946 | 35.39% | 772 | 201 | 3040 |
| Coreidae | Coreinae | Anisoscelini | *Chondrocera* | *laticornis* | 13252384 | 2071848 | 13078 | 5550583 | 424 | 201 | 3056 | 1281 | 47.92% | 667 | 203 | 2880 |
| Coreidae | Coreinae | Anisoscelini | *Holhymenia* | sp. | 23751864 | 3065667 | 11114 | 5316381 | 478 | 180 | 5586 | 925 | 34.61% | 783 | 202 | 4311 |
| Coreidae | Coreinae | Anisoscelini | *Holhymenia* | *tibialis* | 27031152 | 3900460 | 24071 | 10242522 | 426 | 186 | 5329 | 1180 | 44.15% | 682 | 202 | 2733 |
| Coreidae | Coreinae | Anisoscelini | *Leptoglossus* | *clypealis* | 28357464 | 3349221 | 9609 | 4449530 | 463 | 199 | 3866 | 919 | 34.38% | 711 | 202 | 2945 |
| Coreidae | Coreinae | Anisoscelini | *Leptoglossus* | *concolor* | 12280360 | 1387839 | 4591 | 2362520 | 515 | 201 | 3088 | 704 | 26.34% | 696 | 202 | 2829 |
| Coreidae | Coreinae | Anisoscelini | *Leptoglossus* | *gonagra* | 14093344 | 2056938 | 13959 | 5745946 | 412 | 190 | 6330 | 1288 | 48.19% | 528 | 201 | 2612 |
| Coreidae | Coreinae | Anisoscelini | *Leptoglossus* | *phyllopus* | 19413576 | 2226010 | 8031 | 3572496 | 445 | 191 | 2941 | 798 | 29.85% | 604 | 202 | 2941 |
| Coreidae | Coreinae | Anisoscelini | *Leptoscelis* | *tricolor* | 20168192 | 2248553 | 6963 | 3354706 | 482 | 198 | 3099 | 863 | 32.29% | 749 | 202 | 3099 |

| | | | | | | | | | | | | | | | | |
|---|---|---|---|---|---|---|---|---|---|---|---|---|---|---|---|---|
| Coreidae | Coreinae | Anisoscelini | *Narnia* | *femorata* | 16659968 | 2105608 | 7848 | 3664498 | 467 | 190 | 3014 | 938 | 35.09% | 687 | 201 | 2880 |
| Coreidae | Coreinae | Anisoscelini | *Narnia* | *snowi* | 34543752 | 4748128 | 10603 | 5256204 | 496 | 194 | 3515 | 857 | 32.06% | 858 | 201 | 2934 |
| Coreidae | Coreinae | Anisoscelini | *Phthiacnemia* | *picta* | 10478200 | 1060764 | 4179 | 2118784 | 507 | 201 | 19805 | 717 | 26.82% | 561 | 201 | 2634 |
| Coreidae | Coreinae | Chariesterini | *Chariesterus* | *antennator* | 12629816 | 1651822 | 5134 | 2105293 | 410 | 184 | 2649 | 556 | 20.80% | 457 | 202 | 2578 |
| Coreidae | Coreinae | Chariesterini | *Chariesterus* | *armatus* | 34757128 | 5627807 | 24350 | 10745954 | 441 | 180 | 40245 | 1245 | 46.58% | 691 | 202 | 2925 |
| Coreidae | Coreinae | Chariesterini | *Plapigus* | *abdominalis* | 26238680 | 2442042 | 4001 | 1836622 | 459 | 186 | 2931 | 591 | 22.11% | 569 | 202 | 2931 |
| Coreidae | Coreinae | Chelinideini | *Chelinidea* | *tabulata* | 22556288 | 2221185 | 7073 | 3332722 | 471 | 179 | 4619 | 879 | 32.88% | 703 | 202 | 2991 |
| Coreidae | Coreinae | Chelinideini | *Chelinidea* | *vittiger* | 18426880 | 2052586 | 7465 | 3340624 | 448 | 201 | 3539 | 869 | 32.51% | 652 | 201 | 3539 |
| Coreidae | Coreinae | Cloresmini | *Cloresmus* | *antennatus* | 76506720 | 13331549 | 10889 | 4397003 | 404 | 190 | 2918 | 990 | 37.04% | 527 | 201 | 2918 |
| Coreidae | Coreinae | Cloresmini | *Cloresmus* | sp. | 21065080 | 2833560 | 18299 | 8099671 | 443 | 190 | 4481 | 1309 | 48.97% | 691 | 201 | 2739 |
| Coreidae | Coreinae | Cloresmini | *Notobitus* | *meleagris* | 54756520 | 5736099 | 17332 | 6657135 | 384 | 181 | 2977 | 872 | 32.62% | 494 | 201 | 1672 |
| Coreidae | Coreinae | Cloresmini | *Notobitus* | nr *affinis* | 17918224 | 2531762 | 14292 | 6019326 | 421 | 185 | 2939 | 1306 | 48.86% | 635 | 206 | 2939 |
| Coreidae | Coreinae | Cloresmini | *Notobitus* | *sexguttatus* | 31809848 | 5485037 | 24738 | 9999266 | 404 | 177 | 4073 | 1322 | 49.46% | 636 | 203 | 3076 |
| Coreidae | Coreinae | Colpurini | *Homalocolpura* | *parrilloi* | 47417528 | 3768613 | 4922 | 1474106 | 299 | 188 | 1862 | 242 | 9.05% | 259 | 201 | 1188 |
| Coreidae | Coreinae | Colpurini | *Hygia* | sp. | 31714280 | 5256441 | 20990 | 8776844 | 418 | 181 | 4510 | 1380 | 51.63% | 646 | 204 | 2623 |
| Coreidae | Coreinae | Colpurini | *Typhlocolpura* | *chinai* | 21637616 | 1934375 | 3840 | 1774988 | 462 | 198 | 2702 | 558 | 20.88% | 574 | 201 | 2702 |
| Coreidae | Coreinae | Coreini | *Coreus* | *marginatus* | 17361952 | 2855747 | 18057 | 7178637 | 398 | 181 | 2938 | 1159 | 43.36% | 578 | 201 | 2833 |
| Coreidae | Coreinae | Coreini | *Haidara* | *producta* | 13395872 | 1862468 | 13971 | 5997095 | 429 | 186 | 4870 | 1275 | 47.70% | 617 | 202 | 2812 |
| Coreidae | Coreinae | Daladerini | *Dalader* | *planiventris* | 17547888 | 1914916 | 16226 | 6241920 | 385 | 185 | 3313 | 1324 | 49.53% | 545 | 202 | 2557 |
| Coreidae | Coreinae | Daladerini | *Odontocurtus* | *consociatus* | 25154872 | 3865528 | 22366 | 9306231 | 416 | 185 | 4235 | 1372 | 51.33% | 718 | 203 | 3517 |
| Coreidae | Coreinae | Daladerini | *Odontorhopala* | *callosa* | 44219256 | 6630669 | 32721 | 13517625 | 413 | 185 | 8240 | 1470 | 54.99% | 774 | 202 | 3311 |
| Coreidae | Coreinae | Dasynini | *Galaesus* | *hasticornis* | 21715160 | 3225382 | 17693 | 7924886 | 448 | 191 | 3649 | 1410 | 52.75% | 712 | 201 | 3112 |
| Coreidae | Coreinae | Discogastrini | *Savius* | *jurgiosus* | 19564744 | 2532913 | 10585 | 5055550 | 478 | 193 | 3215 | 882 | 33.00% | 771 | 202 | 2898 |
| Coreidae | Coreinae | Gonocerini | *Cletoliturus* | *lituripennis* | 17795080 | 2906396 | 19624 | 8591671 | 438 | 182 | 4549 | 1239 | 46.35% | 692 | 201 | 2735 |
| Coreidae | Coreinae | Gonocerini | *Cletomorpha* | *benita* | 23247312 | 3636413 | 23672 | 9813045 | 415 | 187 | 3010 | 1344 | 50.28% | 624 | 201 | 3010 |
| Coreidae | Coreinae | Gonocerini | *Cletus* | *ochraceus* | 11950416 | 1571786 | 7447 | 3217217 | 432 | 184 | 3025 | 729 | 27.27% | 608 | 201 | 2757 |
| Coreidae | Coreinae | Homoeocerini | *Fracastorius* | *cornutus* | 18136784 | 2570467 | 17274 | 7193126 | 416 | 194 | 3738 | 1292 | 48.34% | 629 | 201 | 2905 |
| Coreidae | Coreinae | Homoeocerini | *Homoeocerus* | *bipustulatus* | 11975096 | 1363942 | 5116 | 2132443 | 417 | 193 | 3749 | 717 | 26.82% | 487 | 201 | 2443 |
| Coreidae | Coreinae | Hypselonotini | *Althos* | *obscurator* | 23044600 | 3188217 | 16184 | 7597953 | 469 | 190 | 5228 | 1370 | 51.25% | 743 | 201 | 2951 |
| Coreidae | Coreinae | Hypselonotini | *Anasa* | *bellator* | 17494152 | 1991901 | 7155 | 3493374 | 488 | 194 | 3651 | 798 | 29.85% | 733 | 201 | 3180 |
| Coreidae | Coreinae | Hypselonotini | *Anasa* | *tristis* | 26646864 | 3876785 | 22884 | 9609936 | 420 | 182 | 5972 | 1390 | 52.00% | 669 | 201 | 2845 |

| | | | | | | | | | | | | | | | | |
|---|---|---|---|---|---|---|---|---|---|---|---|---|---|---|---|---|
| Coreidae | Coreinae | Hypselonotini | *Anasa* | *varicornis* | 29736344 | 2719521 | 4916 | 2227048 | 453 | 192 | 3060 | 646 | 24.17% | 593 | 201 | 3060 |
| Coreidae | Coreinae | Hypselonotini | *Catorhintha* | *guttula* | 16907184 | 1811103 | 5867 | 2980702 | 508 | 201 | 5359 | 804 | 30.08% | 642 | 202 | 2872 |
| Coreidae | Coreinae | Hypselonotini | *Catorhintha* | *texana* | 31445064 | 2423631 | 3951 | 2170013 | 549 | 197 | 3015 | 665 | 24.88% | 646 | 201 | 2882 |
| Coreidae | Coreinae | Hypselonotini | *Cebrenis* | *danieli* | 24995744 | 3320309 | 9938 | 4902308 | 493 | 187 | 3639 | 1027 | 38.42% | 708 | 202 | 2996 |
| Coreidae | Coreinae | Hypselonotini | *Hypselonotus* | *bitrianguliger* | 18789832 | 1953506 | 6527 | 3149896 | 483 | 188 | 6313 | 865 | 32.36% | 672 | 201 | 3005 |
| Coreidae | Coreinae | Hypselonotini | *Hypselonotus* | *fulvus* | 16718296 | 2283263 | 8265 | 4256071 | 515 | 187 | 4594 | 889 | 33.26% | 727 | 204 | 3070 |
| Coreidae | Coreinae | Hypselonotini | *Paryphes* | *pulchellus* | 13304392 | 1884708 | 10787 | 5207546 | 483 | 197 | 4363 | 1269 | 47.47% | 725 | 201 | 2913 |
| Coreidae | Coreinae | Hypselonotini | *Scolopocerus* | *secundarius* | 41109960 | 5962966 | 24573 | 10590577 | 431 | 183 | 4162 | 1251 | 46.80% | 703 | 203 | 2661 |
| Coreidae | Coreinae | Hypselonotini | *Vazquezitocoris* | *repletus* | 16156832 | 2310475 | 12459 | 5642465 | 453 | 183 | 3267 | 1356 | 50.73% | 704 | 201 | 3086 |
| Coreidae | Coreinae | Hypselonotini | *Villasitocoris* | *inconspicuus* | 16751872 | 2525389 | 15824 | 7344793 | 464 | 186 | 4195 | 1240 | 46.39% | 702 | 201 | 3058 |
| Coreidae | Coreinae | Hypselonotini | *Zicca* | *commaculata* | 27665272 | 1981658 | 3750 | 1842088 | 491 | 201 | 3249 | 727 | 27.20% | 550 | 203 | 2610 |
| Coreidae | Coreinae | Hypselonotini | *Zicca* | *rubricator* | 19219952 | 1977073 | 6509 | 3276490 | 503 | 201 | 5336 | 923 | 34.53% | 648 | 201 | 3029 |
| Coreidae | Coreinae | Hypselonotini | *Zicca* | *taeniola* | 54209392 | 5465764 | 10549 | 5480883 | 520 | 179 | 4617 | 1051 | 39.32% | 803 | 202 | 3624 |
| Coreidae | Coreinae | Latimbini | *Latimbus* | *concolor* | 18065568 | 2197106 | 13192 | 6015600 | 456 | 201 | 4642 | 1389 | 51.96% | 690 | 202 | 3068 |
| Coreidae | Coreinae | Latimbini | *Latimbus* | *refulgens* | 14619104 | 1907301 | 13277 | 5989876 | 451 | 193 | 5779 | 1355 | 50.69% | 681 | 202 | 2935 |
| Coreidae | Coreinae | Mictini | *Anoplocnemis* | sp.2 | 32548256 | 2894395 | 5223 | 2360145 | 452 | 188 | 2983 | 657 | 24.58% | 601 | 201 | 2727 |
| Coreidae | Coreinae | Mictini | *Dianomictis* | *expansa* | 10090832 | 1622178 | 14085 | 5603745 | 398 | 185 | 2665 | 1055 | 39.47% | 564 | 201 | 2544 |
| Coreidae | Coreinae | Mictini | *Elasmopoda* | sp. | 28776544 | 2847408 | 9435 | 4601448 | 488 | 191 | 4858 | 816 | 30.53% | 762 | 201 | 2990 |
| Coreidae | Coreinae | Mictini | *Molipteryx* | *lunata* | 20468840 | 3019057 | 21926 | 8124626 | 371 | 181 | 2921 | 1359 | 50.84% | 529 | 201 | 2425 |
| Coreidae | Coreinae | Mictini | *Plectropoda* | *cruciata* | 25782592 | 3678225 | 19248 | 8849062 | 460 | 194 | 3718 | 1370 | 51.25% | 773 | 204 | 3354 |
| Coreidae | Coreinae | Mictini | *Plectropoda* | sp. | 17646344 | 2174687 | 7906 | 3708662 | 469 | 189 | 3184 | 928 | 34.72% | 665 | 201 | 3184 |
| Coreidae | Coreinae | Mictini | *Plectropodoides* | *dallastai* | 17526056 | 2591597 | 14609 | 6998785 | 479 | 193 | 4110 | 1239 | 46.35% | 740 | 201 | 3228 |
| Coreidae | Coreinae | Mictini | *Prionolomia* | *yunnanensis* | 32333088 | 2747554 | 12364 | 5157342 | 417 | 184 | 2946 | 816 | 30.53% | 477 | 201 | 2215 |
| Coreidae | Coreinae | Mictini | *Pternistria* | *bispina* | 17175752 | 2251630 | 15834 | 7199066 | 455 | 187 | 4731 | 1327 | 49.64% | 727 | 203 | 3059 |
| Coreidae | Coreinae | Nematopodini | *Grammopoecilus* | *angustatus* | 21557888 | 3080938 | 16505 | 8047806 | 488 | 183 | 4593 | 1431 | 53.54% | 800 | 205 | 2758 |
| Coreidae | Coreinae | Nematopodini | *Melucha* | *quadrivittis* | 19219872 | 2583836 | 12302 | 5243357 | 426 | 183 | 3158 | 894 | 33.45% | 700 | 202 | 2924 |
| Coreidae | Coreinae | Nematopodini | *Mozena* | *lurida* | 37734920 | 5562609 | 14694 | 7902676 | 538 | 193 | 4911 | 1085 | 40.59% | 935 | 202 | 4911 |
| Coreidae | Coreinae | Nematopodini | *Nematopus* | *lepidus* | 42410856 | 5778383 | 7231 | 2834235 | 392 | 193 | 2606 | 664 | 24.84% | 499 | 201 | 2529 |
| Coreidae | Coreinae | Nematopodini | *Ouranion* | *serrulatus* | 15762120 | 1996195 | 7993 | 3306280 | 414 | 191 | 2961 | 758 | 28.36% | 584 | 202 | 2606 |
| Coreidae | Coreinae | Nematopodini | *Piezogaster* | *calcarator* | 21633944 | 3423138 | 21256 | 9988434 | 470 | 189 | 4990 | 1338 | 50.06% | 790 | 201 | 3085 |
| Coreidae | Coreinae | Nematopodini | *Thasus* | sp. | 33789272 | 4264324 | 15184 | 6920785 | 456 | 178 | 4653 | 975 | 36.48% | 866 | 202 | 3381 |

| | | | | | | | | | | | | | | | | |
|---|---|---|---|---|---|---|---|---|---|---|---|---|---|---|---|---|
| Coreidae | Coreinae | Petascelini | *Petascelis* | *remipes* | 69681960 | 7823970 | 18575 | 8581865 | 462 | 185 | 5952 | 1104 | 41.30% | 867 | 201 | 4042 |
| Coreidae | Coreinae | Phyllomorphini | *Tongorma* | *latreilii* | 42989672 | 7746938 | 61308 | 23175440 | 378 | 180 | 3883 | 1290 | 48.26% | 711 | 203 | 2935 |
| Coreidae | Coreinae | Placoscelini | *Plaxiscelis* | *limbata* | 24768856 | 2844731 | 7688 | 3862816 | 502 | 191 | 5292 | 934 | 34.94% | 755 | 201 | 3137 |
| Coreidae | Coreinae | Placoscelini | *Stenoeurilla* | *mesoamericana* | 17404920 | 1890374 | 7224 | 3465988 | 480 | 195 | 4334 | 840 | 31.43% | 686 | 203 | 3047 |
| Coreidae | Coreinae | Spartocerini | *Sephina* | *geniculata* | 25905048 | 2957702 | 10536 | 5028168 | 477 | 188 | 5901 | 954 | 35.69% | 849 | 204 | 5901 |
| Coreidae | Coreinae | Spartocerini | *Sephina* | *subulata* | 16699200 | 2203106 | 14027 | 6528338 | 465 | 195 | 5420 | 1421 | 53.16% | 700 | 201 | 3633 |
| Coreidae | Coreinae | Spartocerini | *Spartocera* | *fusca* | 33478192 | 3670141 | 13206 | 6385191 | 484 | 201 | 4780 | 1090 | 40.78% | 807 | 201 | 3036 |
| Coreidae | Meropachyinae | Merocorini | *Merocoris* | *curtatus* | 23548936 | 3220266 | 15672 | 7558007 | 482 | 201 | 3633 | 1289 | 48.22% | 775 | 201 | 2955 |
| Coreidae | Meropachyinae | Merocorini | *Merocoris* | *elevatus* | 39274320 | 5288788 | 25274 | 11634030 | 460 | 201 | 7125 | 1352 | 50.58% | 797 | 202 | 3209 |
| Coreidae | Meropachyinae | Merocorini | *Merocoris* | *typhaeus* | 30116232 | 3318342 | 8755 | 4084375 | 467 | 198 | 4391 | 1011 | 37.82% | 661 | 201 | 2860 |

Table S3. Summary data of contigs and ultraconserved element loci successfully obtained from dried museum samples.

| **Family** | **Subfamily** | **Tribe** | **Genus** | **Species** | **Collection year** | **Contigs** | **Total bp** | **Mean contig length** | **Min contig length** | **Max contig length** | **UCE loci** | **Mean UCE length** | **Min UCE length** | **Max UCE length** |
|---|---|---|---|---|---|---|---|---|---|---|---|---|---|---|
| Coreidae | Coreinae | Acanthocerini | *Athaumastus* | *subterlineatus* | 2016 | 17167 | 6838761 | 398 | 194 | 3143 | 645 | 433 | 203 | 1580 |
| Coreidae | Coreinae | Cloresmini | *Notobitus* | *meleagris* | 2015 | 17332 | 6657135 | 384 | 181 | 2977 | 872 | 494 | 201 | 1672 |
| Coreidae | Coreinae | Cloresmini | *Notobitus* | nr *affinis* | 2015 | 14292 | 6019326 | 421 | 185 | 2939 | 1306 | 635 | 206 | 2939 |
| Coreidae | Coreinae | Colpurini | *Homalocolpura* | *parrilloi* | 1946 | 4922 | 1474106 | 299 | 188 | 1862 | 242 | 259 | 201 | 1188 |
| Coreidae | Coreinae | Homoeocerini | *Fracastorius* | *cornutus* | 2015 | 17274 | 7193126 | 416 | 194 | 3738 | 1292 | 629 | 201 | 2905 |

Table S4. Summary of informative sites and number of UCE loci for each dataset.

| Dataset | UCE loci | Total sites | Min number of informative sites in UCE loci | Max number of informative sites in UCE loci | Total Parsimony-informative site (%) | Total Parsimony-uninformative sites (%) | Total Invariant sites (%) |
|---|---|---|---|---|---|---|---|
| 50p, all loci | 1000 | 322084 | 16 | 1079 | 163088 (50.64%) | 21110 (6.55%) | 137886 (42.81%) |
| 50p, 25% most informative loci | 251 | 155772 | 212 | 1079 | 94541 (60.69%) | 11319 (7.27%) | 49912 (32.04%) |
| 70p, all loci | 545 | 202083 | 16 | 1079 | 105112 (52.01%) | 13557 (6.71%) | 83414 (41.28%) |
| 70p, 25% most informative loci | 136 | 100988 | 284 | 1079 | 61463 (60.86%) | 6957 (6.89%) | 32568 (32.25%) |

Table S5. Symmetric distances across all optimal trees (outgroups excluded).

| | ASTRAL-III | | | | RAxML | | | |
|---|---|---|---|---|---|---|---|---|
| | 50p | | 70p | | 50p | | 70p | |
| | 25% most informative | Total evidence | 25% most informative | Total evidence | 25% most informative | Total evidence | 25% most informative | Total evidence |
| **ASTRAL-III, 50p, 25% most informative** | 0 | | | | | | | |
| **ASTRAL-III, 50p, Total evidence** | 8 | 0 | | | | | | |
| **ASTRAL-III, 70p, 25% most informative** | 0 | 8 | 0 | | | | | |
| **ASTRAL-III, 70p, Total evidence** | 2 | 6 | 2 | 0 | | | | |
| **RAxML, 50p, 25% most informative** | 4 | 12 | 4 | 6 | 0 | | | |
| **RAxML, 50p, Total evidence** | 8 | 10 | 8 | 10 | 4 | 0 | | |
| **RAxML, 70p, 25% most informative** | 2 | 10 | 2 | 4 | 2 | 6 | 0 | |
| **RAxML, 70p, Total evidence** | 6 | 8 | 6 | 8 | 8 | 4 | 6 | 0 |

Table S6. Symmetric distances across all trees after branches with less than 50% bootstrap support in optimal trees were collapsed (outgroups excluded).

| | **ASTRAL-III** | | | | **RAxML** | | | |
|---|---|---|---|---|---|---|---|---|
| | **50p** | | **70p** | | **50p** | | **70p** | |
| | **25% most informative** | **Total evidence** | **25% most informative** | **Total evidence** | **25% most informative** | **Total evidence** | **25% most informative** | **Total evidence** |
| **ASTRAL-III, 50p, 25% most informative** | 0 | | | | | | | |
| **ASTRAL-III, 50p, Total evidence** | 8 | 0 | | | | | | |
| **ASTRAL-III, 70p, 25% most informative** | 2 | 8 | 0 | | | | | |
| **ASTRAL-III, 70p, Total evidence** | 4 | 6 | 4 | 0 | | | | |
| **RAxML, 50p, 25% most informative** | 7 | 13 | 5 | 9 | 0 | | | |
| **RAxML, 50p, Total evidence** | 8 | 10 | 8 | 10 | 5 | 0 | | |
| **RAxML, 70p, 25% most informative** | 5 | 11 | 3 | 7 | 2 | 7 | 0 | |
| **RAxML, 70p, Total evidence** | 6 | 8 | 6 | 8 | 9 | 4 | 7 | 0 |

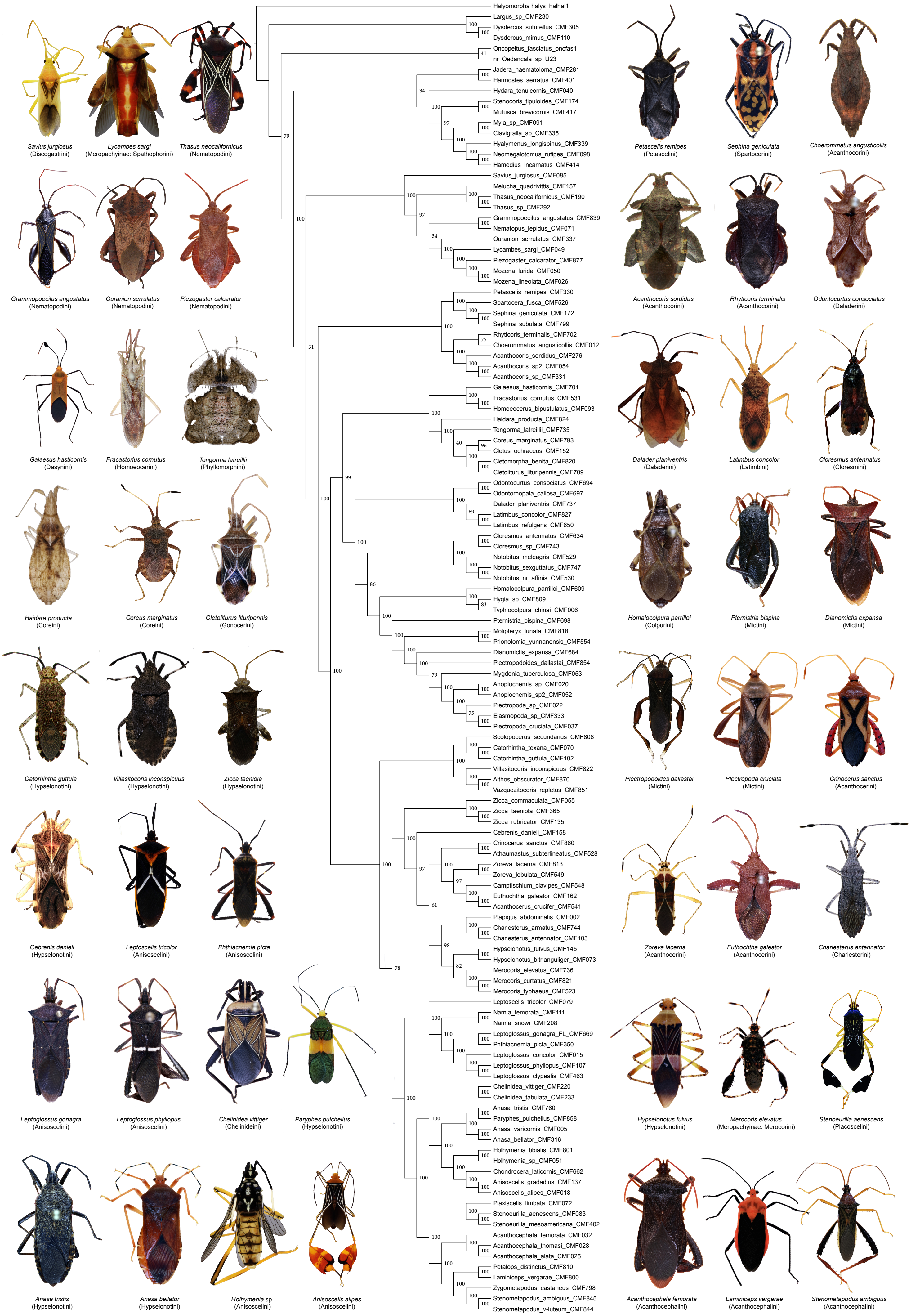

Figure S1. ASTRAL-III species tree generated from all 50p gene trees (i.e., total evidence). Vaues at nodes represent bootstrap support. Dorsal habitus images not to scale.

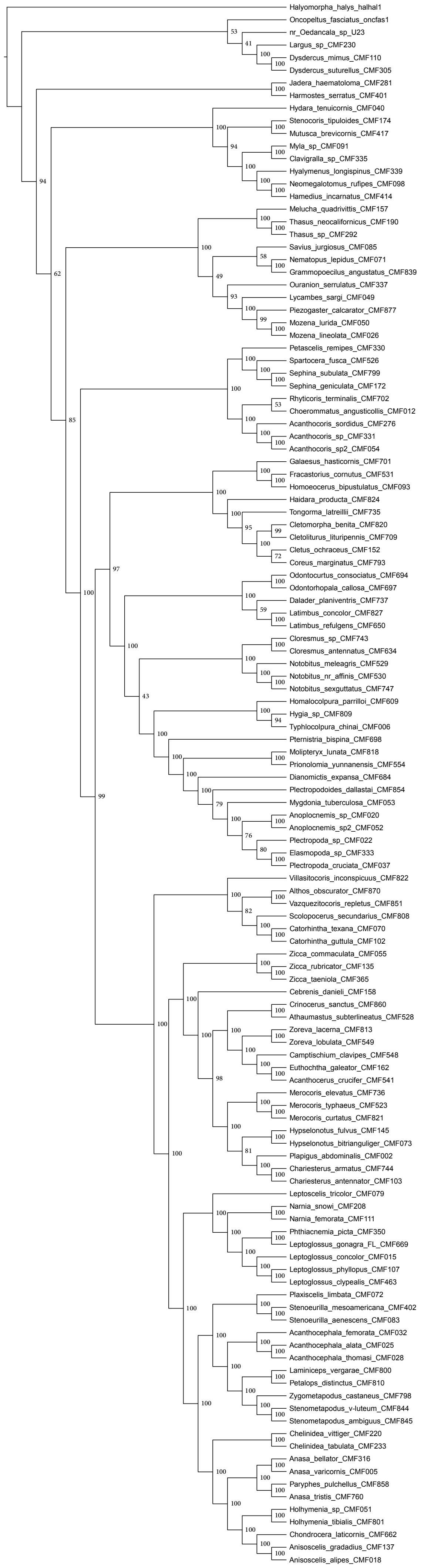

Figure S2. ASTRAL-III species tree generated from the 25% most informative 50% taxon-complete gene trees. Vaues at nodes represent bootstrap support.

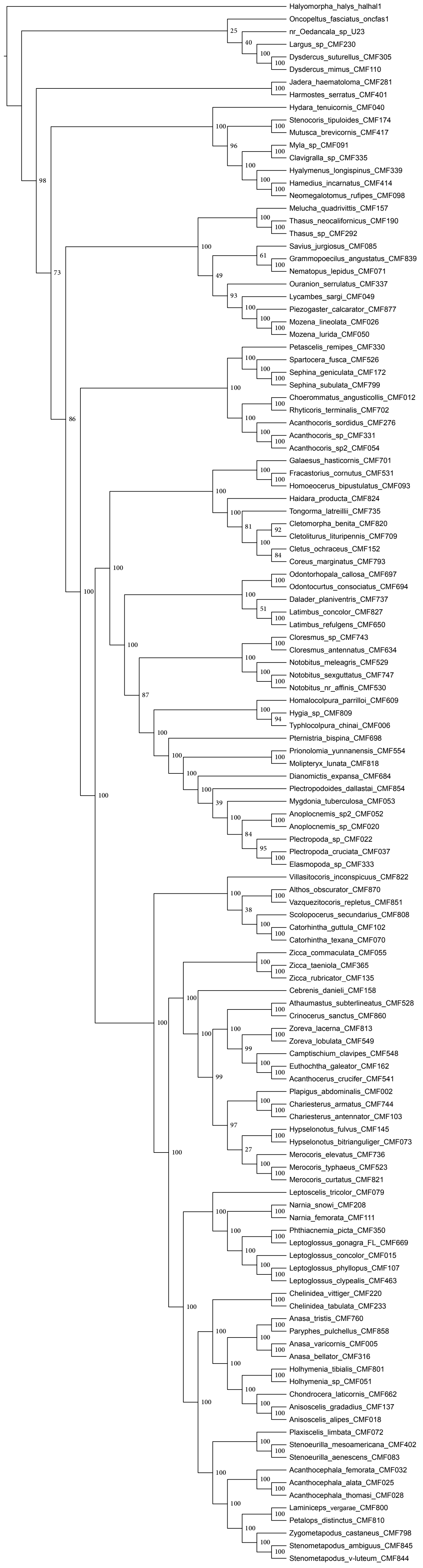

Figure S3. ASTRAL-III species tree generated from all 70p gene trees (i.e., total evidence). Vaues at nodes represent bootstrap support.

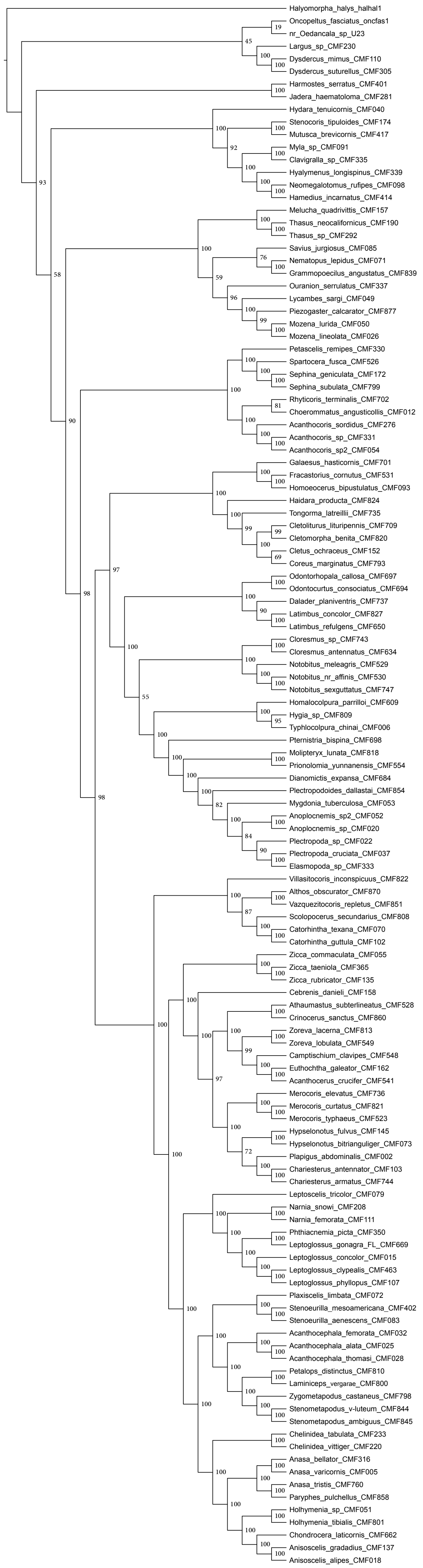

Figure S4. ASTRAL-III species tree generated from the 25% most informative 70p gene trees. Vaues at nodes represent bootstrap support.

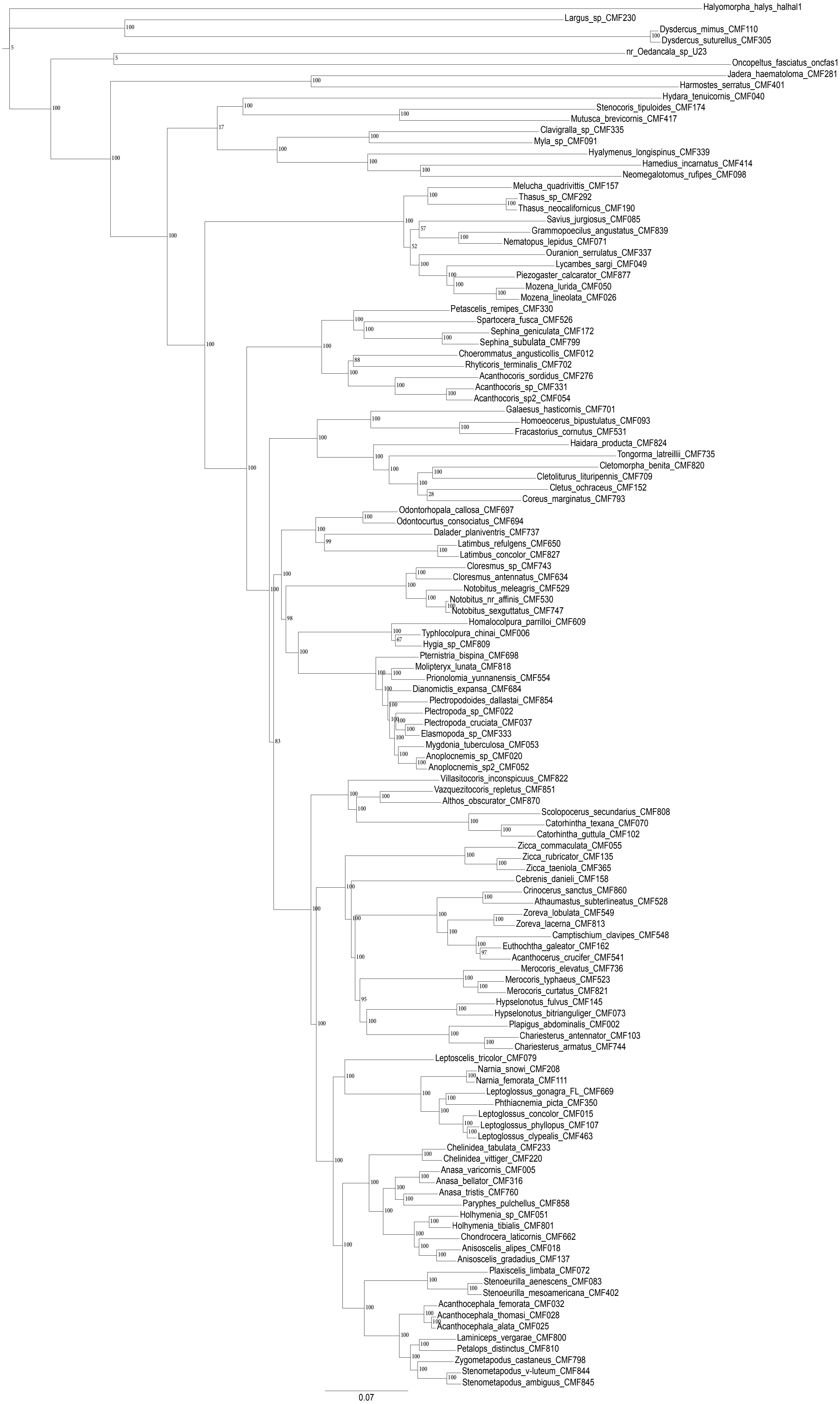

Figure S6. RAxML best tree generated from the 25% most informative 50p gene trees. Values at nodes represent bootstrap support.

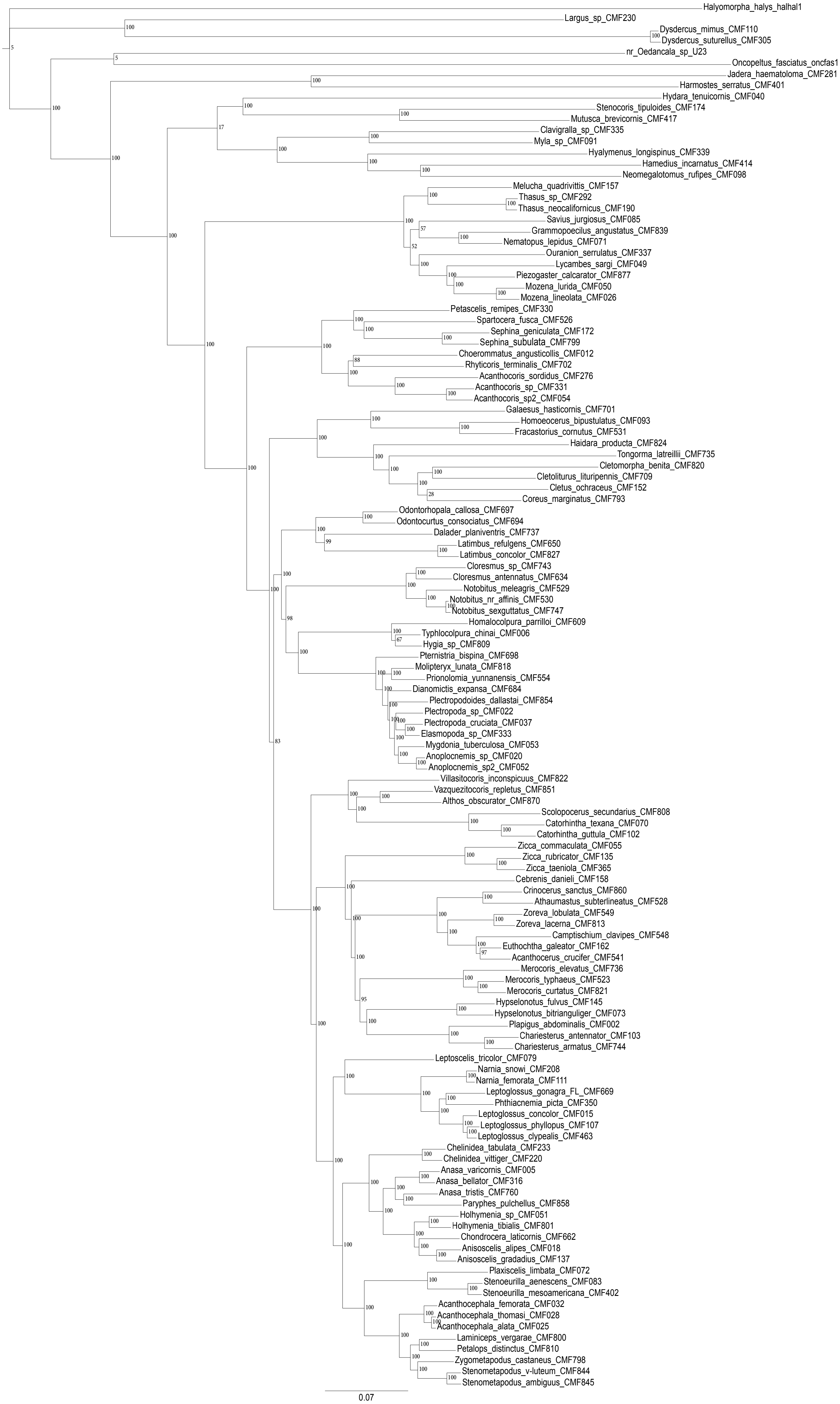

Figure S6. RAxML best tree generated from the 25% most informative 50p gene trees. Values at nodes represent bootstrap support.

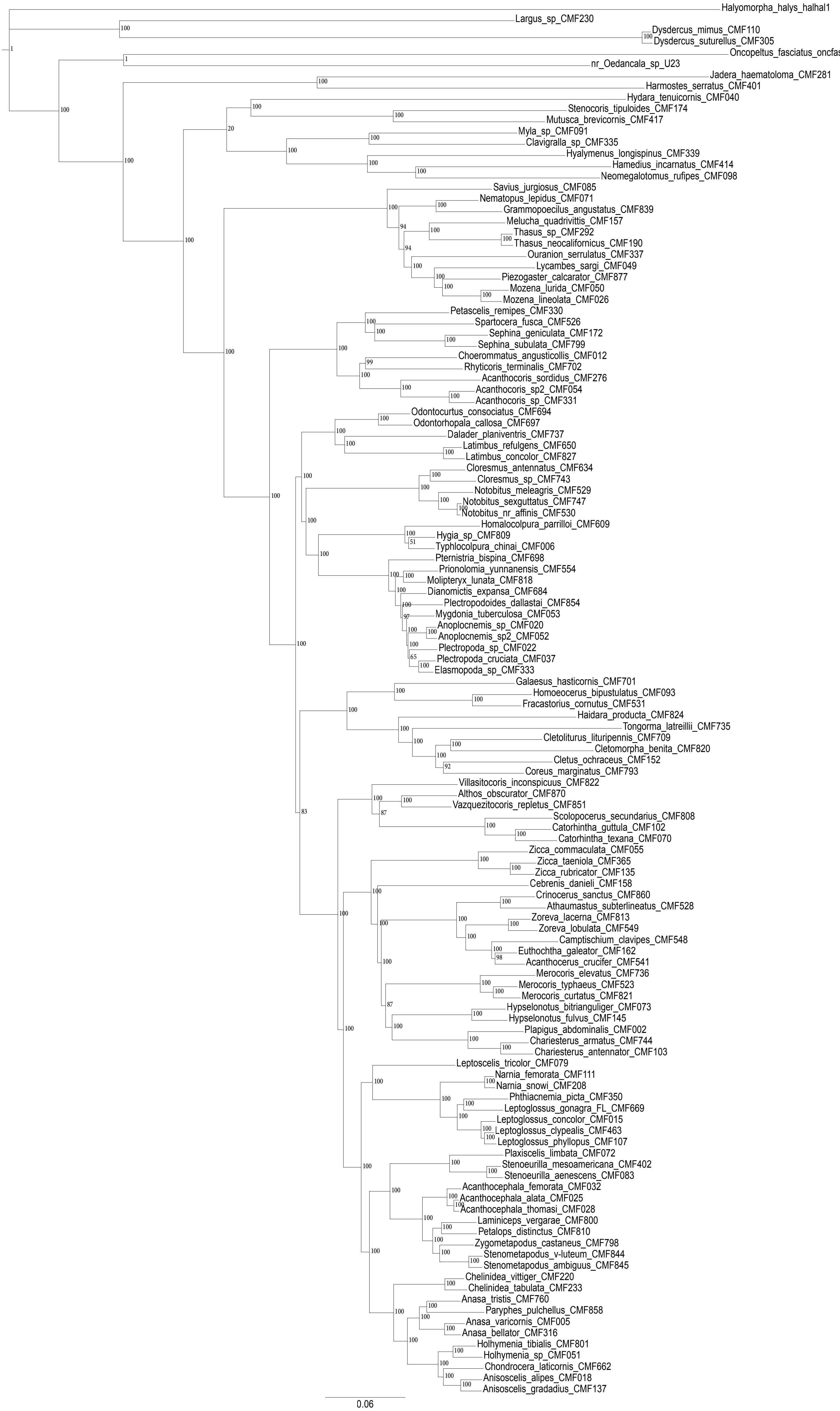

Figure S7. RAxML best tree generated from the 70p gene trees. Values at nodes represent bootstrap support.

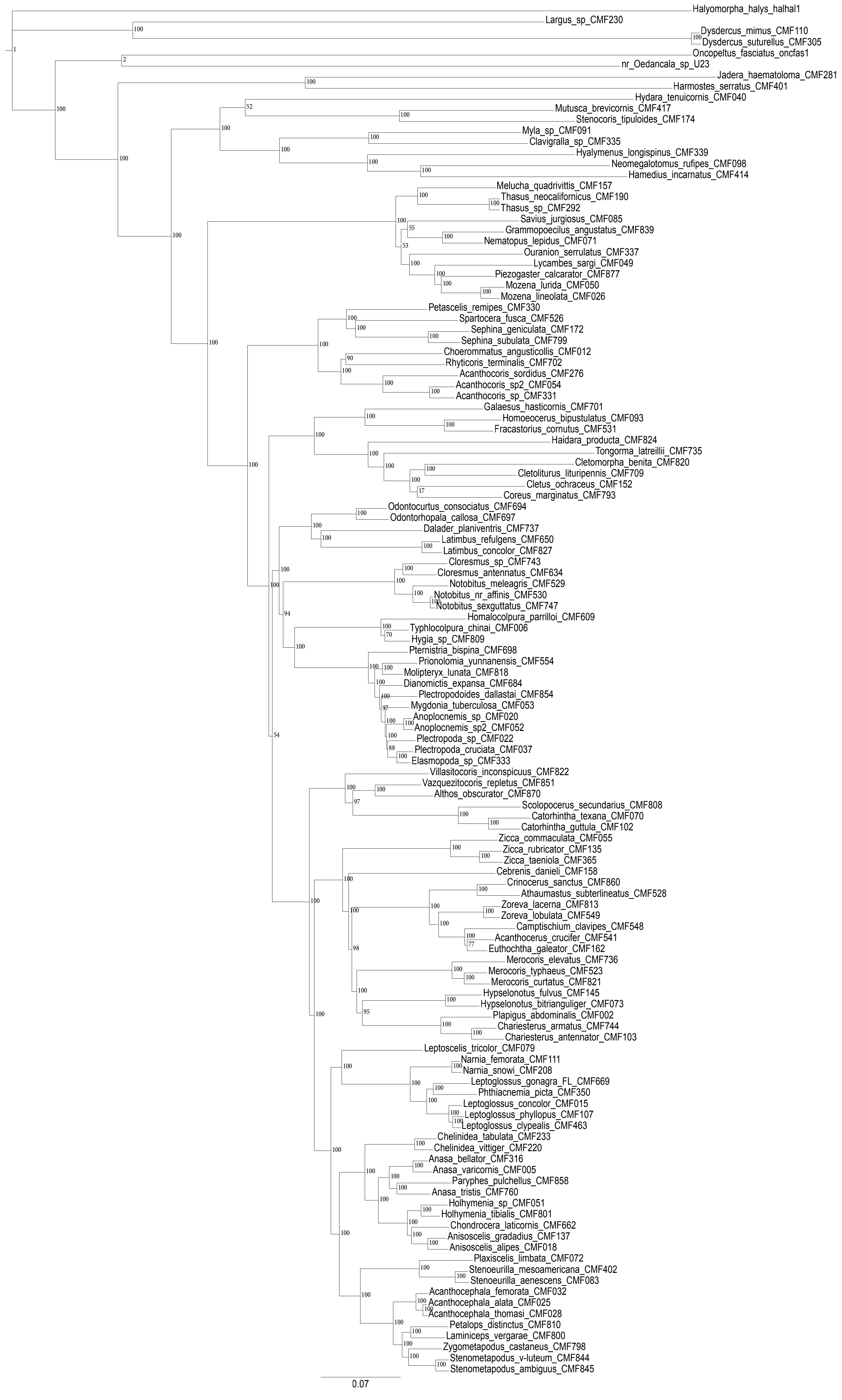

Figure S8. RAxML best tree generated from the 25% most informative 70p gene trees. Values at nodes represent bootstrap support.

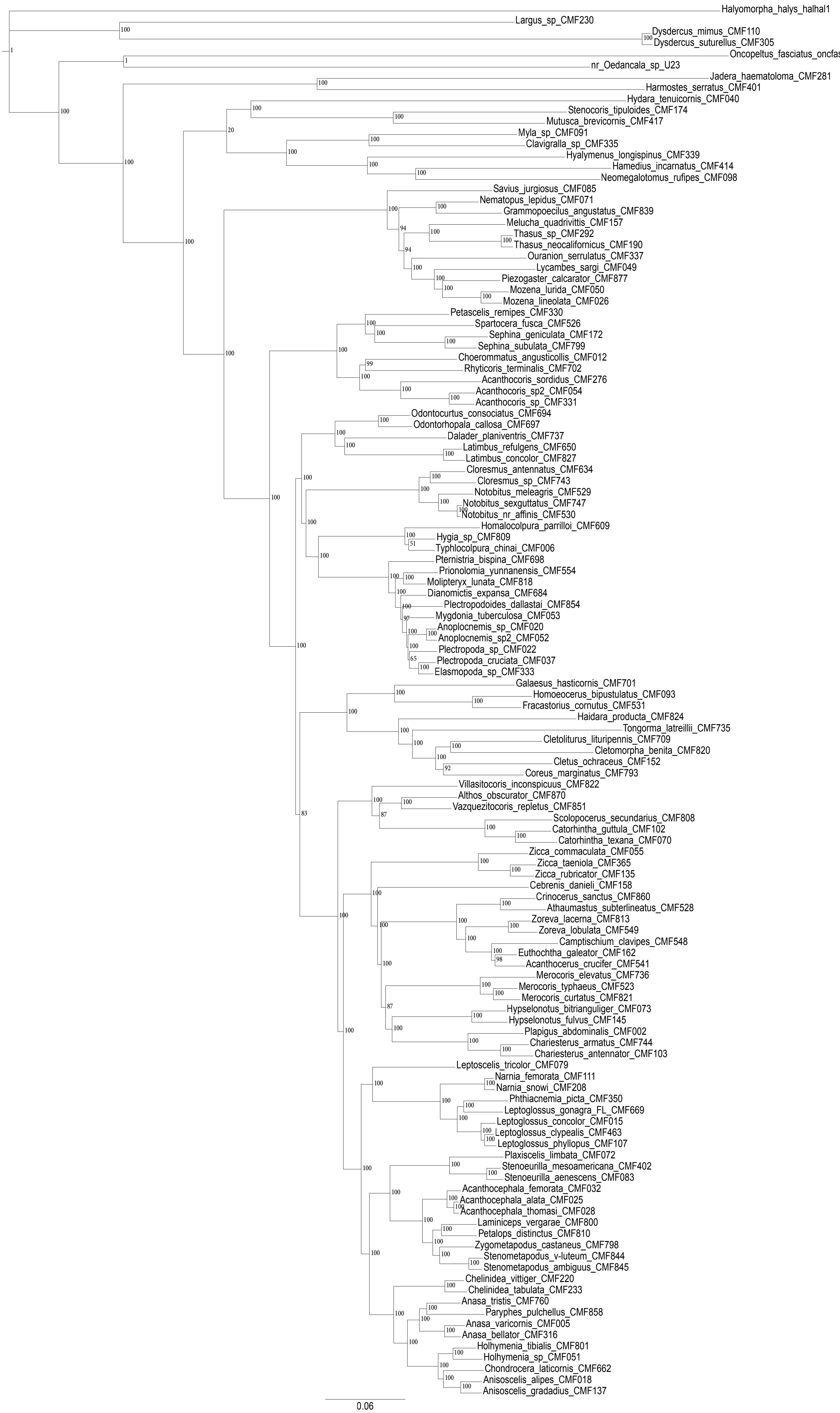

Figure S7. RAxML best tree generated from the 70p gene trees. Values at nodes represent bootstrap support.

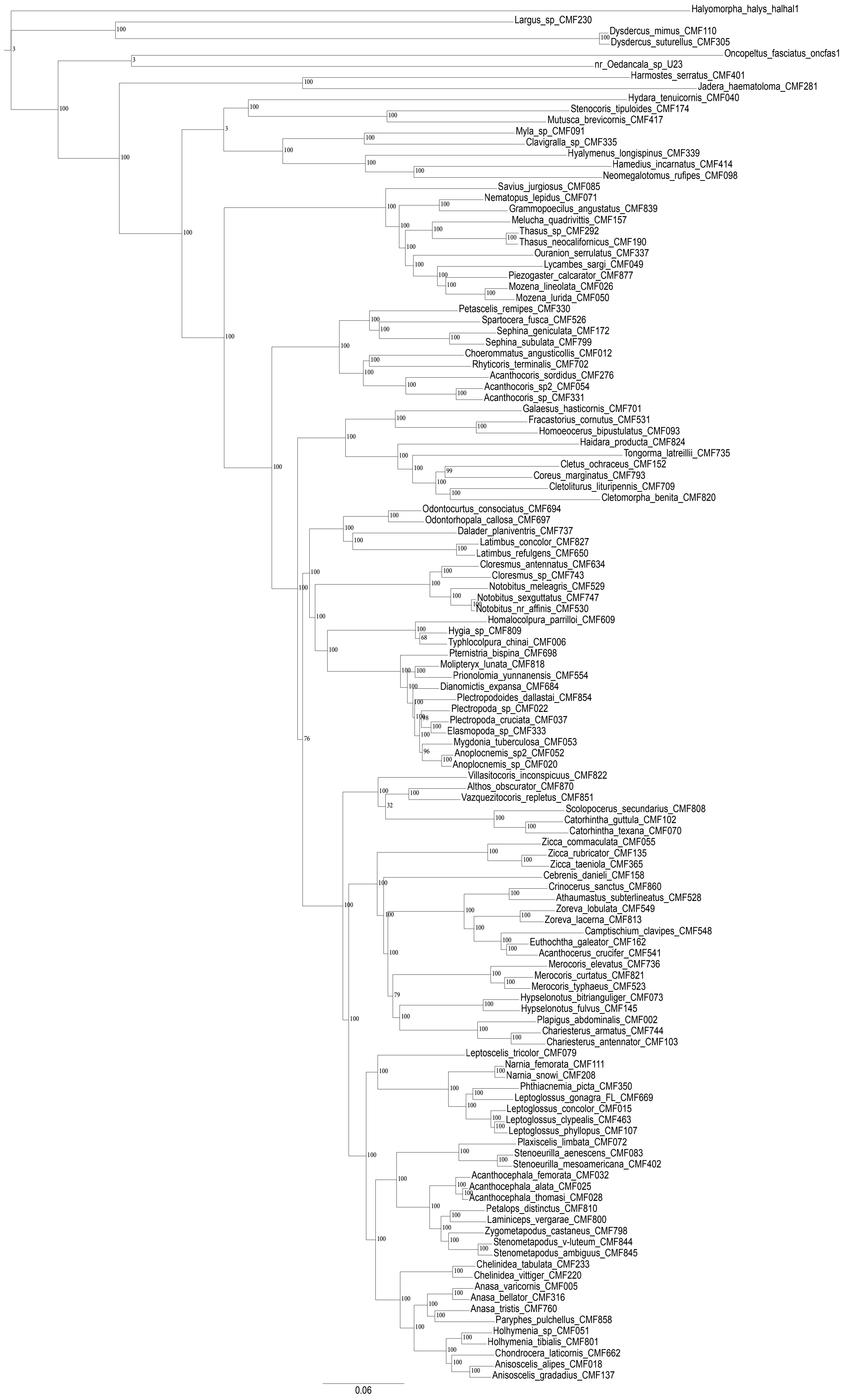

Figure S5. RAxML best tree generated from the 50p gene trees. Values at nodes represent bootstrap support.
